# Salivary Extracellular Vesicle RNA Profiling Reveals Biomarkers for Sjögren’s

**DOI:** 10.1101/2025.08.05.668729

**Authors:** Sudipto K. Chakrabortty, Shuran Xing, Allan George, Benjamin Sawicki, Steven Lang, Sinead Nguyen, T. Jeffrey Cole, Emily Mitsock, Christian Ray, Driss Zoukhri, Mabi Singh, Loukas Chatzis, Andreas Goules, Maria-Ioanna Saridaki, Sivakumar Gowrisankar, Athanasios G Tzioufas, Athena Papas, Johan Skog

**Affiliations:** Division of Oral Medicine, Tufts School of Dental Medicine, Boston MA, USA; Exosome Diagnostics, a Bio-Techne brand, Waltham, MA, USA; Research Institute for Systemic Autoimmune Diseases, Athens, Greece; Department of Pathophysiology, School of Medicine, National and Kapodistrian University of Athens, Athens, Greece; Laboratory of Immunobiology, Center for Clinical, Experimental Surgery and Translational Research, Biomedical Research Foundation of the Academy of Athens, Athens, Greece

## Abstract

Sjögren’s is a chronic autoimmune disease affecting exocrine glands and is subclassified into SSA-positive (SSA+) and SSA-negative (SSA-) subtypes, with a complex diagnostic journey and an average diagnostic delay of almost 4 years. While SSA+ cases can be detected via serological testing, current assays lack specificity. For SSA-patients, no non-invasive diagnostic tools exist, and diagnosis often requires invasive lip biopsy. A saliva-based liquid biopsy capable of diagnosing both subtypes is therefore of high clinical interest. However, saliva poses challenges due to its abundant oral microbiome, which complicates unbiased biomarker discovery. In this study, we present a novel RNA sequencing workflow that efficiently depletes microbial content, enabling deep profiling of long RNAs within salivary extracellular vesicles (EVs). This approach identified both known and novel RNA biomarkers capable of diagnosing SSA+ and SSA-subtypes with high sensitivity and specificity. Moreover, we uncovered distinct RNA signatures that allow molecular stratification of Sjögren’s subtypes. Pathway analysis in SSA+ cases revealed enrichment of immune and glandular pathways consistent with prior tissue-based studies, supporting the utility of salivary EVs as a non-invasive surrogate for tissue biopsy. Importantly, our data provides new molecular insights into the under-characterized SSA-subtype, laying the foundation for future mechanistic studies and facilitating their broader inclusion in clinical trials.

## Introduction

Sjögren’s Disease (SjD) is a chronic autoimmune disorder characterized by lymphocytic infiltration of the exocrine glands, leading to hallmark symptoms such as chronic dryness in the mouth (xerostomia) and eyes (xerophthalmia) [1, 2]. Beyond glandular manifestations, SjD frequently involves systemic features, including fatigue, musculoskeletal pain, gastrointestinal symptoms such as GERD, and neurological involvement. In clinical trials, the EULAR Sjögren’s syndrome disease activity index (ESSDAI) additionally assesses involvement in 12 domains (cutaneous, respiratory, renal, articular, muscular, peripheral nervous system (PNS), central nervous system (CNS), hematological, glandular, constitutional, lymphadenopathic, biological) [3]. Patients with SjD are also at increased risk for thyroid disease and mucosa-associated lymphoid tissue (MALT) lymphomas, underscoring the systemic and potentially serious nature of the disease [4, 5]. SjD is currently subclassified serologically based on the presence (SSA+) or absence (SSA-) of antibodies against the Ro antigen [6–8]. The Ro antigen itself is a complex of proteins and small RNAs (Y RNAs), with two main components: Ro60 (also known as TROVE2), which associates with Y RNAs and participates in RNA quality control, and Ro52 (also known as TRIM21), which plays a role in type I interferon (IFN) responses [9–13]. While the presence of SSA/Ro autoantibodies facilitates diagnosis of the SSA+ subtype, these antibodies are also seen in other autoimmune diseases, reducing their specificity for SjD [7, 14, 15]. In addition to established serological markers, McCoy et al. identified novel antibodies – targeting DTD2 and RESF1 – that aid in the diagnosis of anti-SSA-SjD [16]. Diagnosing SjD remains challenging, with an average diagnostic delay of up to 6 years, especially in SSA-patients who lack reliable non-invasive biomarkers [17–19]. These patients must often undergo invasive minor salivary gland biopsy (MSGB) for confirmation—a procedure associated with risks such as pain, nerve damage, and permanent sensory deficits [20–22]. Furthermore, even among SSA+ patients, clinical heterogeneity and varying treatment responses highlight the need for molecular tools to better stratify patients beyond serology alone [23].

Extracellular vesicles (EVs), including exosomes, are nanoscale lipid bilayer-bound vesicles (30–150 nm) secreted by virtually all cells. EVs carry a complex cargo of biomolecules, including messenger RNAs (mRNAs), long non-coding RNAs (lncRNAs), microRNAs (miRNAs), Y-RNAs, tRNAs, proteins, lipids, and DNA [24, 25]. These vesicles mediate intercellular communication and reflect the molecular state of their cell of origin, making them an attractive target for biomarker discovery in diseases like SjD [26–28]. Saliva is a highly accessible biofluid enriched with EVs, offering a compelling platform for non-invasive diagnostics [29, 30]. While several studies have profiled small RNAs in salivary EVs [31–37], comprehensive profiling of *long* RNAs from salivary EVs has remained elusive [38, 39]. This is largely due to the overwhelming presence of microbial RNA in saliva, which hampers detection of the human transcriptome and limits the depth of sequencing coverage of human target genes [40, 41].

In this study, we developed a novel transcriptomic workflow that overcomes the microbial RNA barrier and enables ultra-deep sequencing of human long RNAs from salivary EVs. By applying this technology to patients across the SjD spectrum, including SSA+, SSA-, healthy controls, and Sicca-positive/SjD-negative individuals—we aimed to identify salivary EV RNA biomarkers capable of subclassifying SjD patients by molecular phenotype and uncover previously uncharacterized transcriptomic signatures unique to the SSA-subtype.

Our findings reveal that salivary EV long RNA profiles harbor distinct and reproducible gene expression signatures that differentiate SjD subtypes and illuminate underlying molecular heterogeneity. This work represents a major step toward developing a non-invasive, molecularly informed classification system for SjD. In particular, the ability to identify and stratify SSA-patients—who currently lack a non-invasive diagnostic option—marks a significant clinical advance. Ultimately, EV transcriptome profiling holds promise not only for early diagnosis but also for patient stratification and targeted therapeutic development in SjD.

## Results

Development of saliva as a platform for liquid biopsy has been limited due to lack of standardized methods for collection & pre-processing of saliva [31, 40]. At the outset, we began by establishing a robust saliva collection and pre-processing workflow for isolation of EVs and RNA downstream (see methods). As part of this study, de-identified saliva samples were obtained from a total of 103 individuals, comprising of 24 SSA+ patients, 47 SSA-patients, 16 from healthy individuals, and 16 Sicca patients (Figure 1b). Additionally, to ensure specificity of the biomarker signature, we also profiled 5 Rheumatoid arthritis and 5 Lupus patients. The clinical metadata for all donors is summarized in Supplementary Table 1 and 2. A representative Bioanalyzer RNA Pico profile of saliva EV RNA is shown in Supplementary Figure 1.

**Figure 1:**
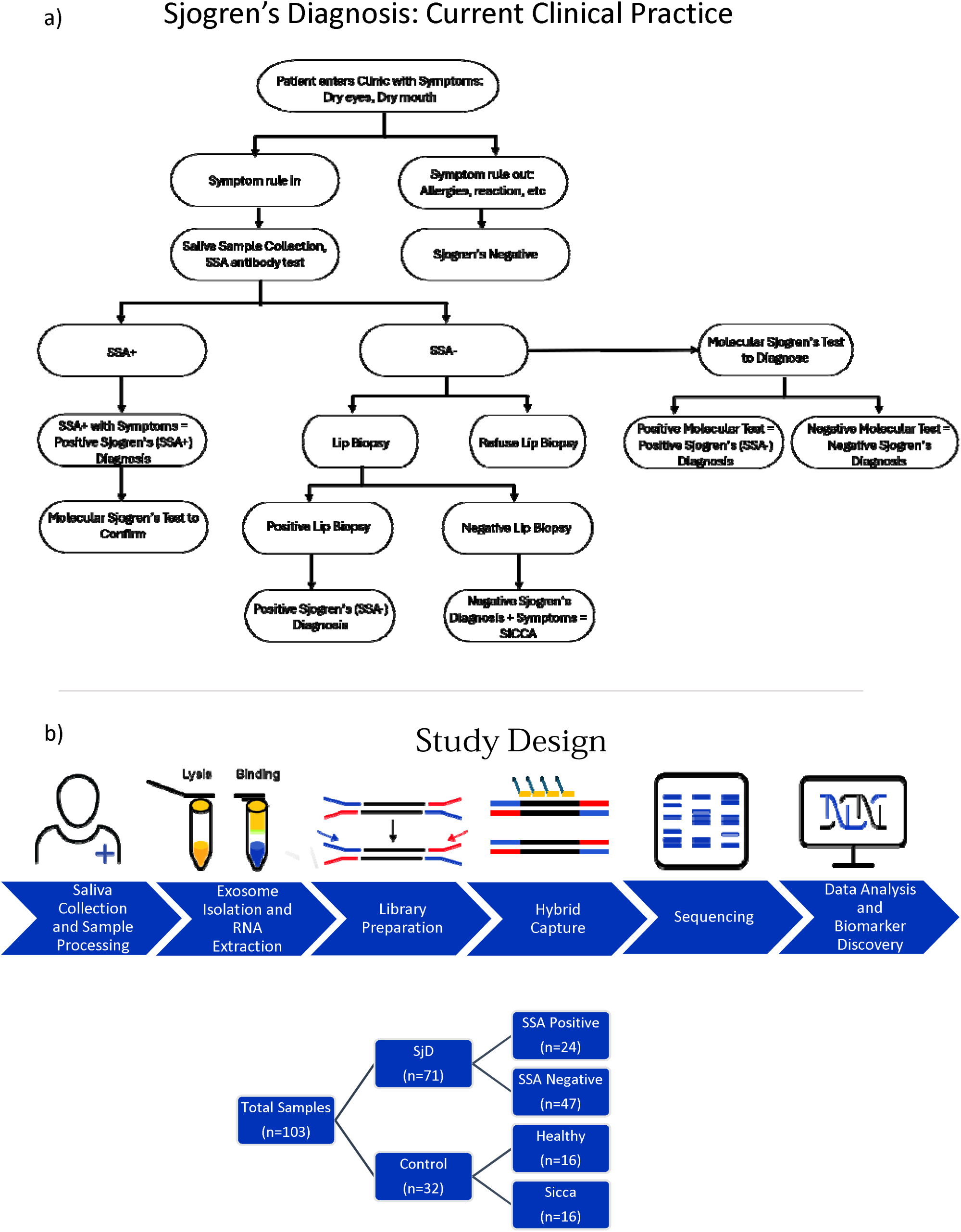
Sjogren’s Syndrome Clinical Diagnosis and Study Workflow Overview a) Flowchart of current clinical practice for the diagnosis of Sjogren’s Syndrome. b) Study design and sample summary of Sjogren’s study.

Previous efforts to profile saliva EV RNAs have reported that the overwhelming majority of RNASeq reads cannot be aligned to the human genome, suggesting the microbial origin of these reads [40, 42]. Yeri et. al. and Rozowsky et. al. showed only a third (34% +-8.7%) of the RNASeq reads could be aligned to the human genome, while nearly half of the reads (46% +-11.9%) remain unaligned to the human genome [40, 42] . To address these challenges, we developed a novel RNASeq workflow that combines ultra-sensitive library construction strategies with hybrid capture technology. In a stark contrast to previous results, the majority of RNASeq reads (79% +-11.7%) obtained from our workflow could be aligned to the human genome resulted, while only 1% of the reads could not be aligned to the human genome (Figure 2a). The remarkably low proportion of unmapped reads in our data underscores the highly efficient depletion of the salivary microbiome derived signal achieved by our hybrid capture based RNASeq workflow. Consequentially, the pronounced enrichment of human EV RNA signal achieved by our RNASeq workflow enabled us to profile human salivary EV transcriptome at an unprecedented level. Over 55% of reads were mapped uniquely to the human genome by STAR [43], while approx. another 15 - 40% of reads were multi-mappers (Supplementary Figure 2a). When the genomic origin of uniquely mapped reads was investigated, we observed over 80% of reads mapped to the exons, while approximately 5.7% and 1.9% of reads mapped to introns and intergenic regions, respectively (Supplementary Figure 2b). Biotype distribution of exonic reads revealed over 90% of reads mapping to protein coding genes, followed by approximately 3.2% and 1.0% of reads mapping to long non-coding RNAs and ERCC spike-ins, respectively (Figure 2b). On average, over 12,500 protein coding genes and over 1600 long non-coding RNAs were detected from salivary EVs, at a minimum detection threshold of 0.01 transcripts per million (TPM) (Figure 2c). As expected, tissue deconvolution analysis of saliva EV RNAs revealed enrichment of signal originating from salivary glands and acinar cells of saliva, thus contrasting with abundance of platelet and erythrocytic signal in plasma EVs (Figure 2d, Supplementary Fig. 2c).

**Figure 2:**
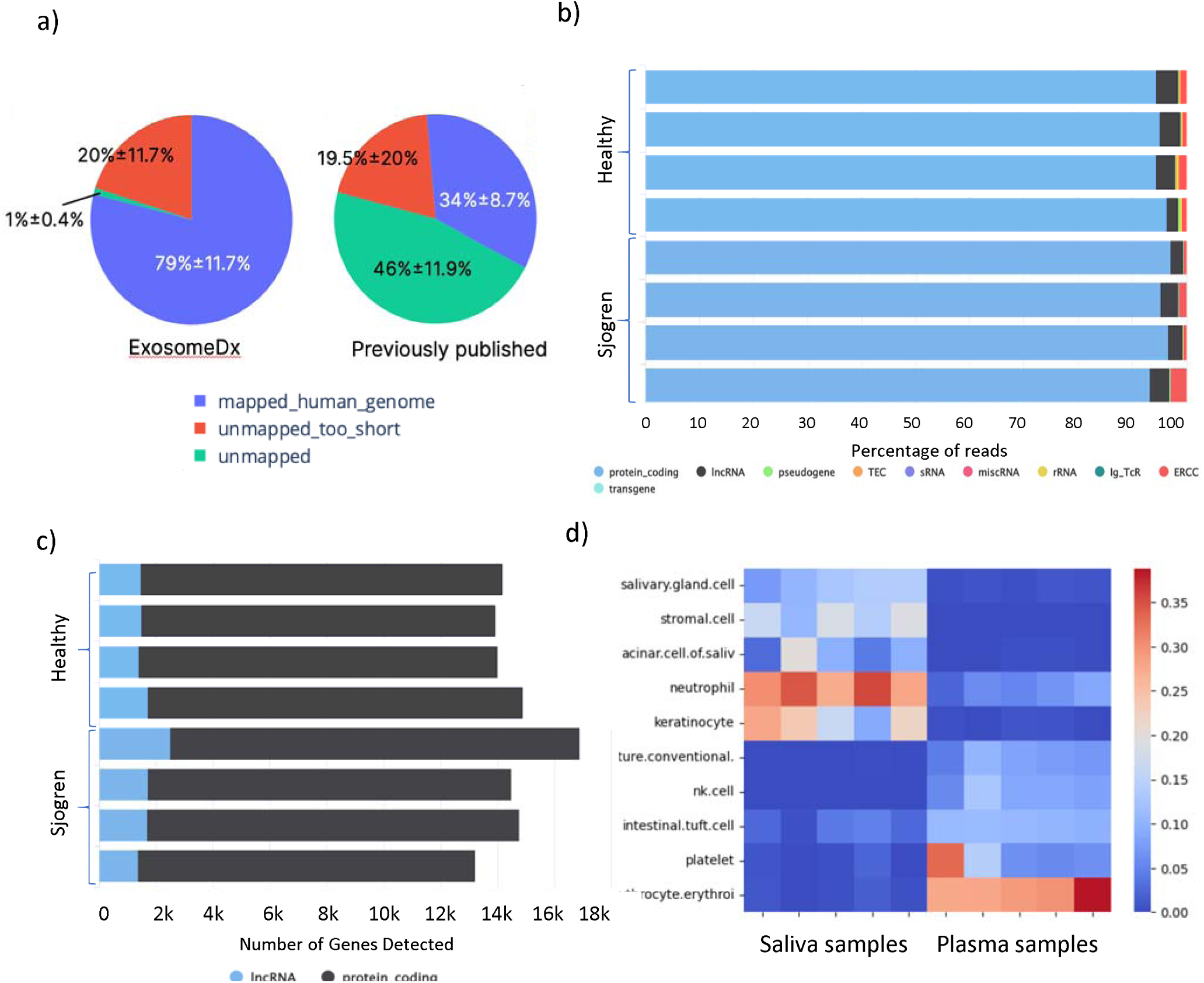
Performance of ExosomeDx’s Proprietary Saliva EV RNA Sequencing Workflow a) Comparison of saliva EV RNA mapping statistics using ExosomeDx’s saliva RNASeq workflow versus published methods. ExosomeDx’s workflow shows a significant increase in the percentage of reads mapping to the human genome. b) Biotype distribution of transcriptomic reads for representative saliva samples. The percentage of mapped reads corresponding to biotypes is shown on the x-axis. Major represented biotypes are protein-coding genes (blue), lncRNAs (black), rRNAs (yellow), and ERCC (red). c) Gene detection of representative saliva samples. The x-axis shows the number of protein coding genes (black) and lncRNAs (blue) detected at a threshold of greater than 0.01 transcripts per million (TPM). d) Deconvolution heatmap contrasting the top five tissue contributors in saliva samples with plasma samples. Five representative samples were used for each biofluid. Salivary gland and erythrocytes showed highest contribution in saliva and plasma, respectively.

Armed with this workflow, we set out to first identify biomarkers of SSA+ subtype. We profiled salivary EV RNAs from 24 SSA+ patients and 32 controls (16 healthy and 16 Sicca). Differential Expression (DEx) analysis between SSA+ patient samples vs healthy and Sicca controls identified 108 genes upregulated and 71 genes downregulated in SSA+ patients by at least two-fold (p adj. < 0.05) (Figure 3a, Supplementary Tables 3&4). Pathway analysis of the genes upregulated in SSA+ using MSigDB database revealed significant enrichment of several Hallmark inflammatory and interferon response pathways (Figure 3c) [44, 45]. Pathway analysis of the upregulated genes in SSA+ using KEGG & Reactome database further corroborated the enrichment of multiple interferon response pathways (Supplementary Figure 3). All three interferons, i.e., interferon alpha, beta and gamma response pathways were found to be significantly upregulated in SSA+ patients, when compared with controls. In particular, the Interferon Alpha response pathway was found to be the most significantly upregulated in SSA+ samples, with nearly 200-fold enrichment over expected (FDR<0.001) (Supplementary Figure 4) [46]. Not only all 31 genes in this pathway were detectable in our SSA+ EV samples, as many as 22/31 genes in this pathway were significantly upregulated in SSA+ samples, underscoring the level of enrichment of this pathway [46]. Additionally, and in complete agreement with literature and the established definition of the SSA+ condition, tissue deconvolution analysis of the salivary EV RNAs, demonstrated a reduced contribution from salivary glands in SSA+ when compared with controls (Figure 3b) [6–8].

**Figure 3:**
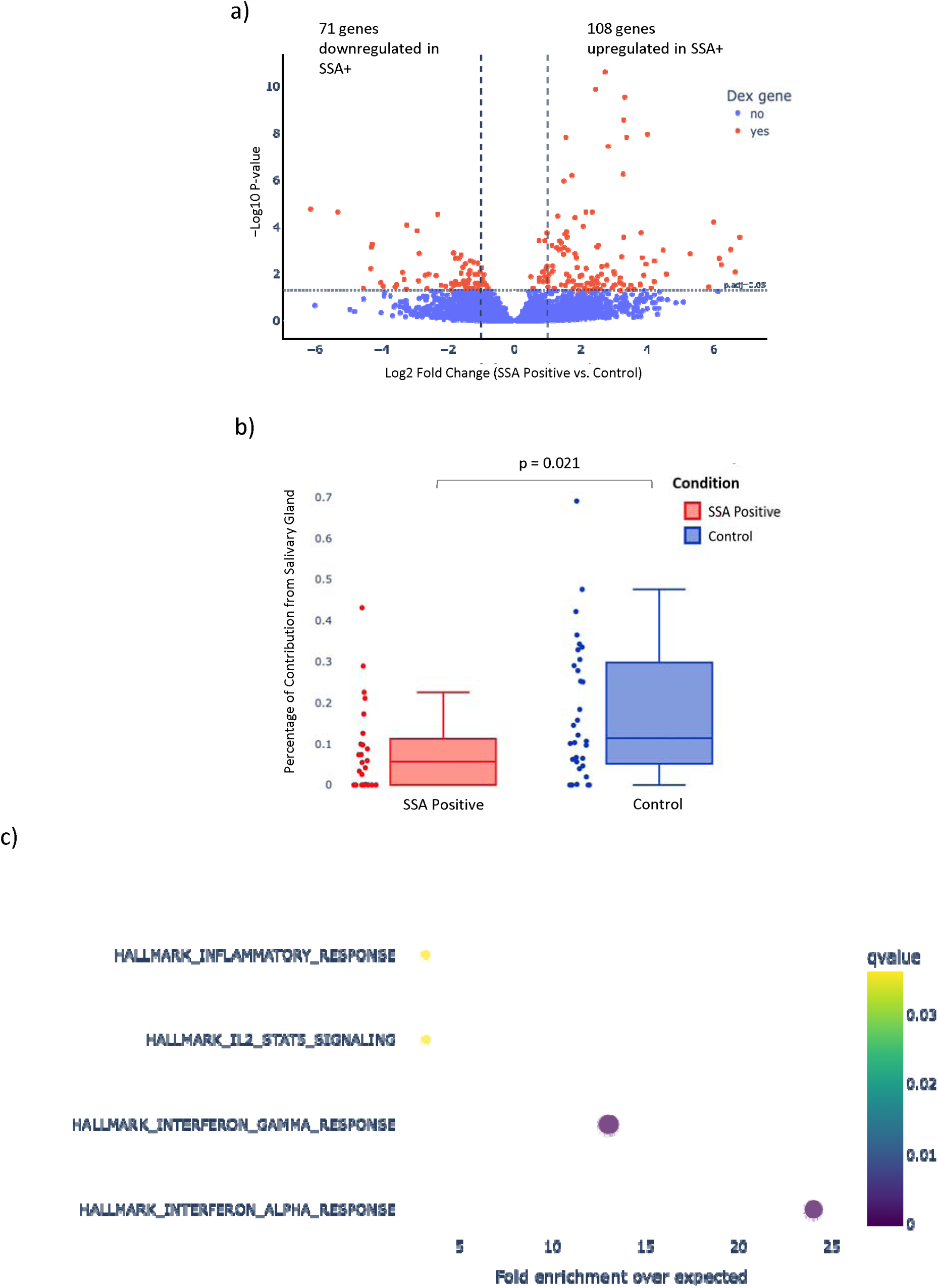
Differential Expression and Pathway Analysis of SSA+ vs Control Samples a) Volcano plot of differentially expressed genes between SSA+ Sjogren’s samples vs Control samples (Healthy and Sicca samples). Genes in red have p-values of ≤ 0.05 and a log2 Fold Change > |1|, while genes in blue have p-values > 0.05 or a log2 Fold Change −1 < x < 1. b) Tissue deconvolution analysis using GTex database as reference shows significant (Mann-Whitney U, p<0.05) downregulation of salivary gland composition in SSA+ Sjogren samples in comparison to control samples. c) Pathway analysis of genes upregulated in SSA+ samples compared to control based on Hallmark gene set database. The x-axis represents fold enrichment over expected, while bubble color denotes the adjusted q-value. Notably, Hallmark interferon-α and interferon-γ response were significantly upregulated in SSA+ samples over control.

Next, we aimed to perform biomarker discovery and identify a saliva EV RNA based signature that is capable of diagnosing SSA+ from healthy and Sicca controls with high sensitivity and specificity. Using a nested CV approach as described in the methods, we were able to achieve an AUC of 0.82 on test data and estimated the sensitivity of 73%, at the specificity of 82% in clinical practices (Figure 4a&c). Additionally, when only Sicca samples were used as controls, the estimated sensitivity was 72% with 82% specificity (data not shown). Features that are highly conserved and selected in more than 30% of all folds in the nested CV are reported in Figure 4d. All feature selected genes were found to be upregulated in SSA+. Principal component analysis based on this signature showed separation of SSA+ samples from healthy and Sicca control samples (Figure 4b). Notably, when extended to include patients with other autoimmune conditions with overlapping symptoms such as rheumatoid arthritis (RA) and lupus, the gene signature demonstrated reasonable separation of SSA+ patients from these autoimmune conditions, suggesting that the gene signature identified may be disease specific (Supplementary Figure 11). This is particularly significant given that traditional SSA antibody testing often lacks the specificity to distinguish Sjögren’s from RA or Lupus [16].

**Figure 4:**
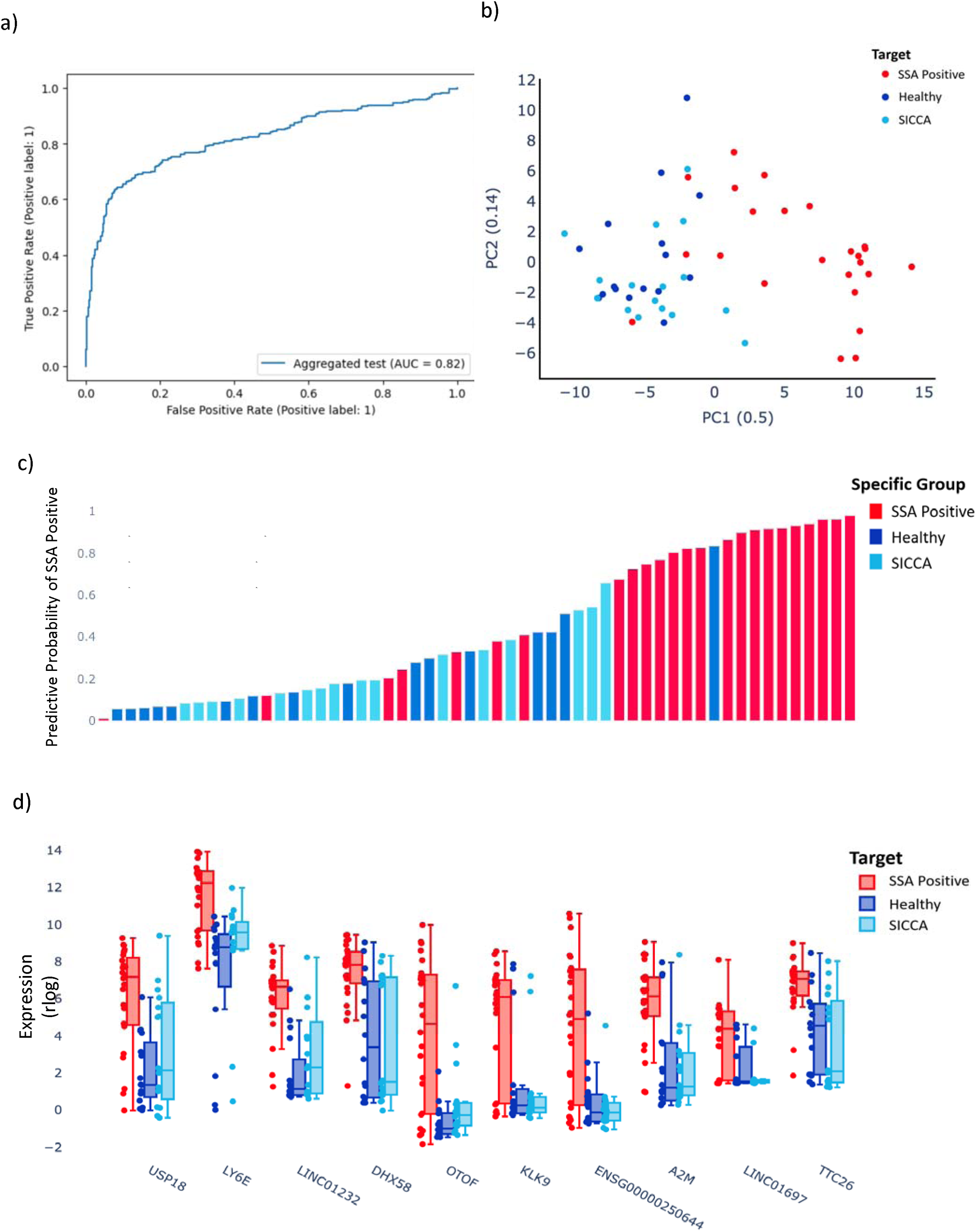
Sjogren’s SSA+ gene signature a) ROC curve of SSA+ gene signature, demonstrating an AUC of 0.82 for classifying SSA+ samples. b) Principal component analysis (PCA) of SSA+ (red), healthy (blue) and Sicca (light blue) samples based on SSA+ 10 gene signature showed separation of SSA+ from controls. c) Waterfall plot demonstrating the performance of SSA+ gene signature. SSA+ samples are shown in red, healthy samples in blue and Sicca samples in light blue. d) Box plots representing the expression levels (TPM) of the 10-gene SSA+ signature across Healthy (blue), Sicca (light blue), and SSA+ samples (red).

In contrast to SSA+, molecular mechanisms underlying SSA-subtype are still poorly understood and require diagnosis with an invasive lip biopsy, resulting in limited enrollment in ongoing clinical trials [12]. To better understand the molecular mechanisms underlying SSA-subtype, we profiled saliva EV transcriptome from 47 SSA-patients and 32 controls, including 16 healthy and 16 Sicca patients by our RNASeq workflow. DEx analysis between SSA-patients vs healthy and Sicca controls identified 308 genes being upregulated and 204 genes downregulated in SSA-, when compared with healthy and Sicca controls (Figure 5a). Principal component analysis using these DEx genes showed clear separation between most SSA-samples and controls (Figure 5b). Interestingly, pathway analysis of genes upregulated in SSA-revealed dysregulation of synapse-associated and sensory signaling pathways, among others (Figure 5c). This is further corroborated by GO cell-of-origin pathway analysis of upregulated genes, which revealed an increased contribution from olfactory neuroepithelium, retinal microglia, midbrain neurotypes and major microglia types (Supplementary Fig. 5). In contrast, pathways related to desmosome and various binding pathways, including alpha catenin binding, clathrin binding, cadherin binding, vesicles coat and granules related pathways, among others were found to be downregulated in SSA-samples, when compared with controls (Supplementary Fig.7).

**Figure 5:**
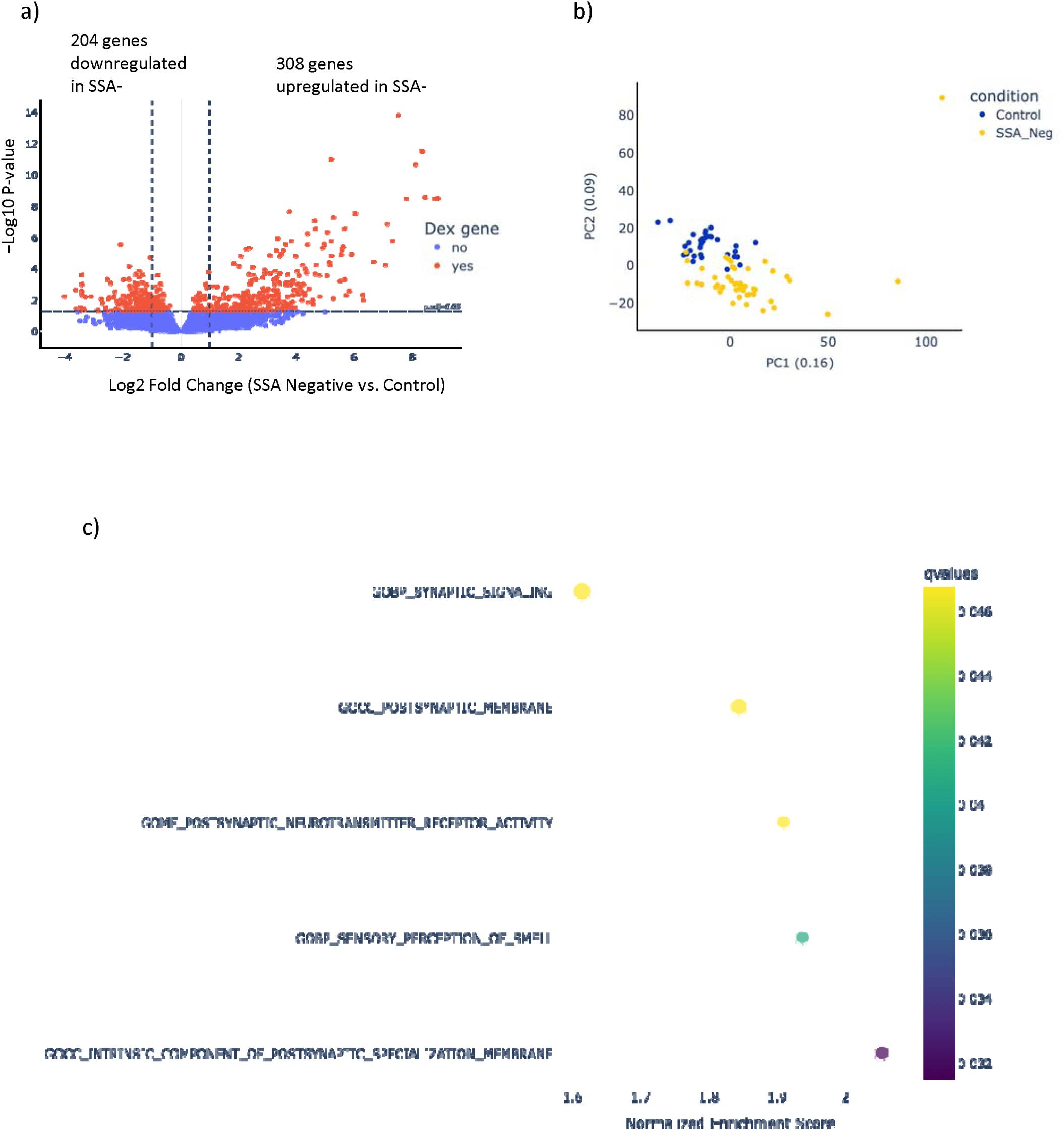
Differential Expression and Pathway Analysis of SSA-vs Control Samples a) Volcano plot showing differentially expressed genes between SSA-samples and Control samples (Healthy and Sicca samples). Genes shown in red have p-values of ≤ 0.05 and a log2 Fold Change > |1|, while genes shown in blue have p-values > 0.05 or a log2 Fold Change −1 < x < 1. b) Principal component analysis between SSA-(red) and Control (blue) samples. SSA- and Control samples show distinct clustering. c) Pathway analysis of differentially expressed genes upregulated in SSA-samples compared to control samples (Healthy and Sicca) using a Gene Set Enrichment Analysis (GSEA), based on the GO database. The x-axis represents normalized enrichment score, while bubble color denotes the adjusted q-value.

Next, we performed biomarker discovery to identify a saliva EV RNA signature to diagnose SSA-patients from healthy and Sicca controls. Using the nested CV approach once again, we were able to achieve an AUC of 0.83 on test data only and estimated the sensitivity of 74%, at the specificity of 82% in clinical practices (Figure 6a&c). Additionally, when only Sicca samples were used as controls, the estimated sensitivity was 72% with 82% specificity (data not shown). Features that are highly conserved and selected in more than 30% of all folds in the nested CV are reported in Figure 6d. Most selected genes are upregulated in SSA-with one exception, TEX41. Principal component analysis based on this signature showed separation of SSA-samples from healthy and Sicca control samples (Figure 6b). When RA and Lupus samples were included, moderate separation was observed, with the majority of SSA-samples forming a loosely defined cluster distinct from most RA and SLE samples (Supplementary Figure 12). This may suggest that the SSA-gene signature captures some disease-specific features despite partial overlap.

**Figure 6:**
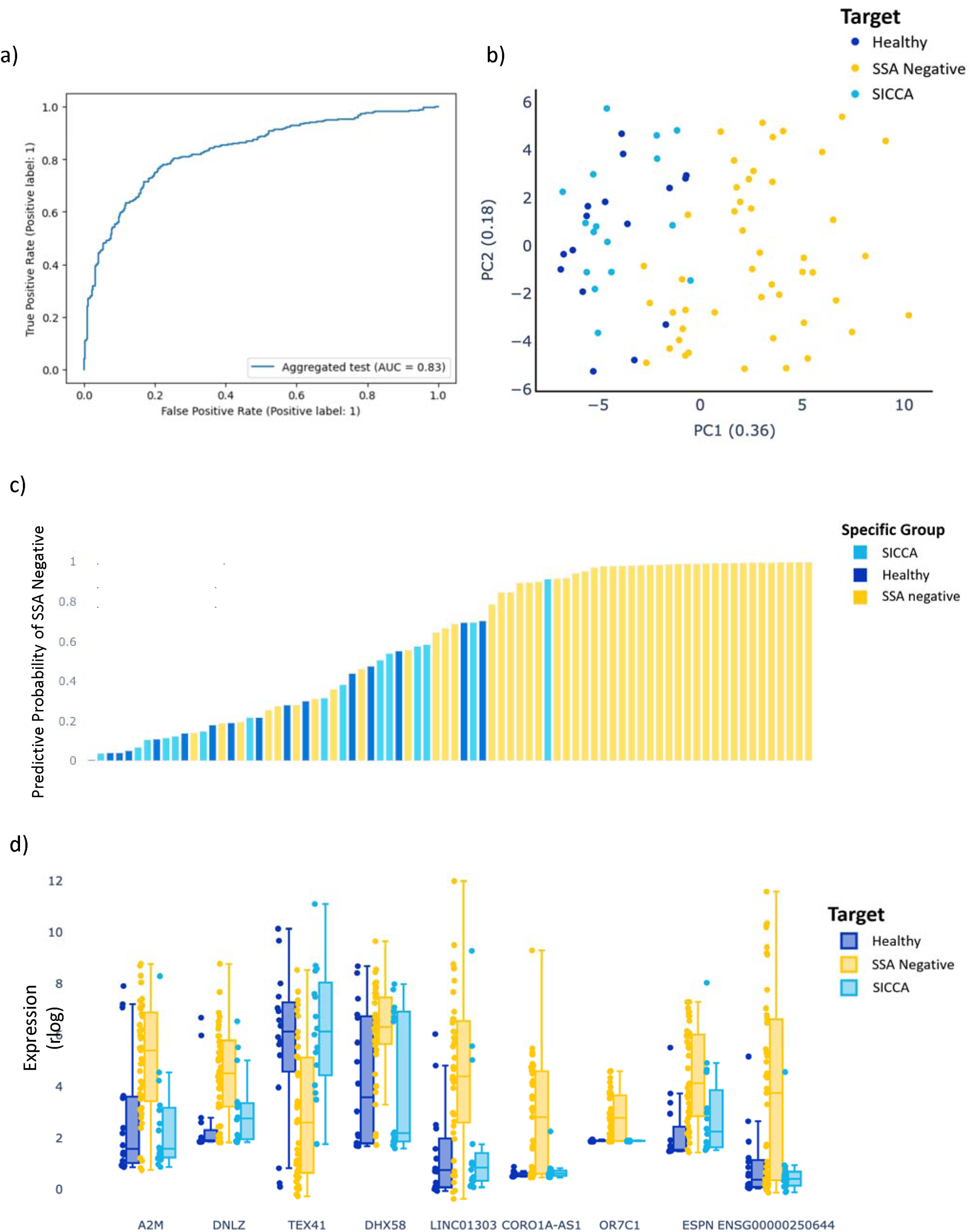
Sjogren’s SSA-Signature Performance a) ROC curve of our SSA-gene signature showing an AUC = 0.83. This 7-gene signature distinguishes SSA-Sjogren’s patients from healthy and Sicca patients with 83% accuracy. b) Principal component analysis (PCA) of SSA-(yellow), healthy (blue) and Sicca (light blue) based on our SSA-gene signature. c) Waterfall plot showing the performance of our SSA-gene signature on all samples. SSA-samples are in yellow, healthy samples are in blue, and Sicca samples are in light blue. d) Comparison of expression levels for the 9 genes which make up our SSA-signature across SSA-(yellow), healthy (blue), and Sicca (light blue) samples.

Finally, we examined the ability of saliva EV RNA to perform a non-invasive, molecular subtyping of SjD. To that end, we performed DEx analysis between SSA+ vs SSA-patients, which identified a total of 324 differentially expressed genes, 84 upregulated in SSA+ and 240 upregulated in SSA-subtype (Figure 7a, Supplementary Table 7&8). Principal component analysis using these DEx genes showed clear separation between the two subtypes (Figure 7b). Pathway analysis of genes upregulated in SSA+ revealed significant enrichment of inflammatory response, interferon-alpha and gamma response pathways in SSA+ subtype (Supplementary Figure 8). In contrast, pathway analysis of genes upregulated in SSA-pointed to the enrichment of multiple pathways related to synapse assembly, organization and localization, gliogenesis, nervous system development and granule membrane, among others (Supplementary Figure 9). Cell of origin analysis of genes upregulated in SSA-further revealed increased contribution from eye microglia, retina microglia, other microglia types and retina fibroblast, among others (Supplementary Figure 11). When taken together, these results suggest that completely different underlying molecular mechanisms might be at play in the two SjD subtypes.

**Figure 7:**
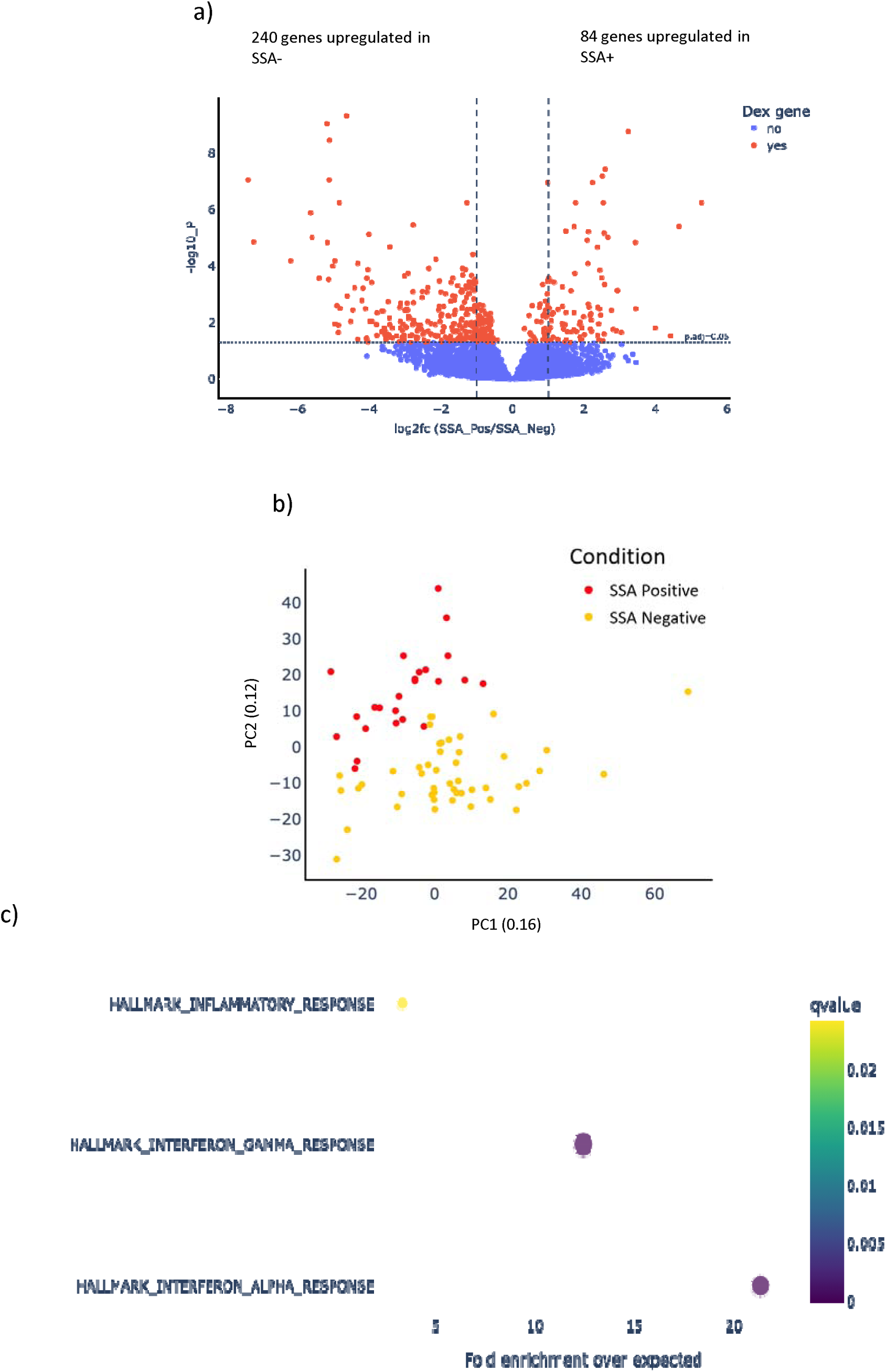
Differential Expression of SSA-Samples a) Volcano plot showing differentially expressed genes between SSA+ and SSA-samples. Genes shown in red have p-values of ≤ 0.05 and a log2 Fold Change > |1|, while genes shown in blue have p-values > 0.05 or a log2 Fold Change −1 < x < 1. b) Principal component analysis between SSA+ (red) and SSA-(yellow) samples. SSA+ and SSA-samples show separation. c) Pathway analysis of differentially expressed genes upregulated in SSA+ samples compared to SSA-samples using an Over Representation Analysis (ORA), based on the Hallmark gene set database. The x-axis represents fold enrichment over expected, while bubble color denotes the adjusted q-value.

## Discussion

The clinical landscape of Sjögren’s Disease has several gaps and is challenged by lack of non-invasive diagnostic solutions. On one hand, lack of any noninvasive diagnostic option forces SSA-patients to resort to painful lip biopsy procedure, on the other hand, SSA+ autoantibody test lacks specificity for SjD. Saliva based liquid biopsy has the potential to revolutionize the treatment landscape of SjD. However, lack of standardized processing guidelines has severely limited the emergence of saliva based liquid biopsy. In this study, we tie together several novel approaches (see methods) that overcome the known challenges in saliva processing and serve as a template for standardized collection and pre-processing guidelines for the future. These innovations include use of stimulated saliva, RNase treatment of saliva to reduce RNA fragmentation, dilution of saliva with PBS to reduce viscosity for easier filtration, filtering saliva with 0.8um to remove contaminants and debris, and use of state-of-the-art technology for isolation of EVs from saliva, among others.

Saliva based liquid biopsy has been severely limited due to challenges in EV RNA profiling. Challenges include low quantity and fragmented size distribution profile of saliva EV RNA, lack of sensitive library construction workflows for long RNA profiling, as well as the abundance of oral microbiome in saliva. The novel workflow described in this study not only demonstrated highly efficient depletion of the human microbiome but also enabled highly sensitive profiling of saliva EV long RNAs at an unprecedented depth. To the best of our knowledge, this data represents the most comprehensive profiling of the landscape of long RNA transcriptome from salivary EVs reported to date.

In this study, biological pathway & GO analysis of differentially expressed genes dysregulated in SSA+ have repeatedly pointed to the involvement of multiple Interferon response pathways. This is important, as Interferon response pathways have been previously reported to be dysregulated in SSA+ salivary gland tissue RNASeq studies and has been heavily implicated in SSA+ pathophysiology [47, 48]. While genome-wide association studies (GWAS) have identified fewer loci for SjD compared to other autoimmune conditions, recent work by Lessard et al. has uncovered 22 genome-wide significant loci with functional relevance in both immune and glandular epithelial cells [49] . The limited number of identified loci reflects the clinical and molecular heterogeneity of SjD, which necessitates larger, well-characterized cohorts for biomarker discovery. Ongoing studies continue to explore how genetic, environmental, and epigenetic factors interact to drive disease onset and progression.

Aquaporin gene family members have been suggested to be biomarkers for SSA+ in literature, specifically Aquaporin 5 (AQP5) [50–52]. Results obtained in this study did not find AQP5 to be significantly differentially expressed when comparing either SSA+ or SSA-groups to the combined control cohort of healthy and Sicca. Interestingly, when the control group was stratified into healthy and Sicca subgroups, AQP5 expression was significantly downregulated in both SSA+ and SSA-samples relative to the Sicca group, while the downregulation was found to be not significant when compared to healthy controls (Supplementary Figure 13). This may suggest that AQP5 downregulation may be more specific to SjD pathology than to Sicca symptoms alone. The lack of significant downregulation compared to healthy samples may be attributed to differential sample processing, different molecular readout technologies being utilized as well as relatively smaller sample size of this study. Deconvolution analysis of saliva EV protein coding genes in this study demonstrated reduced contribution from salivary gland cells in SSA+ patients when compared with healthy controls. This is critical, as SjD itself is defined by the killing of the salivary gland cells by the body’s own immune system [53]. Taken together, these data highlight the ability of salivary EVs to reliably capture the transcriptomic landscape and dysregulated pathways in salivary gland tissue, demonstrating its potential to be an excellent, non-invasive surrogate of tissue biopsy.

Previous studies have focused on dysregulated miRNAs in salivary epithelial and mesenchymal cells. Conversely, exogenous EVs have been found to enter salivary gland cells and can affect Wnt signaling, affecting cell activity and secretory function [54]. Alevizos et al. first described differences in minor salivary gland miRNA in SjD vs healthy individuals, and suggested that microRNA may be protective of epithelial cells [55]. Additionally, it has been shown by D. Iwakiri et. al., that Epstein–Barr virus-microRNAs interfere with calcium signaling, leading to salivary gland hypofunction [56]. Overexpression of miR-181a and miR-16 in SjD patients has been associated with saliva hypofunction downregulation of let-7b miRNA increased expression of miR-181a, miR-200b and miR-223 miRNAs [56]. In a recent review article SjD patients had 94 miRNAs differentially expressed regulating inflammatory and neurological pathways that affect secretions compared to controls [57].

In this study, we identified several robust gene signatures in saliva EVs as potential biomarkers of SSA+. Five of these genes have known relevance in SjD, lending further credence to these biomarkers, while the other five are putative novel biomarkers that require further exploration. For example, LY6E, an interferon induced gene, has been previously identified as a biomarker for SSA+ [58]. Not only it is known to be upregulated in SjD, but it has also been identified as a “hub” gene, playing a central role in Sjögren’s molecular network [59]. Similarly, the gene A2M has also been identified as a potential therapeutic target [60]. Not only A2M has been previously reported to be dysregulated in SjD, like LY6E, it has also been suggested as a “hub” gene for SjD.[60, 61]. The gene OTOF encodes otoferlin, a protein involved in neurotransmitter release from inner hair cells also has been previously reported to have altered gene expression level and has been identified as a potential biomarker for SjD [59, 62]. USP18, a negative regulator of type I IFN signaling, has been suggested to play a significant role in dysregulation of IFN pathways in SjD and is currently being explored as a potential biomarker [63–65]. While DHX58 has not been directly implicated in SjD, DHX58 plays a role in innate immune response and is involved in interferon pathway, dysregulation of which has been heavily implicated in SjD [66, 67].

Similarly, while KLK9 has not been directly implicated in SjD, the broader family of KLK genes and their role in autoimmune disease, e.g., is being explored [68, 69].

While very limited knowledge currently exists regarding the molecular mechanisms underlying SSA-subtype, and represents a major novelty of this study, it is worth noting that the several of the pathways that have been identified in this study have previously been implicated in SjD. For example, pathway analysis of DEx genes in SSA-in this study pointed to the dysregulation of pathways related to synaptic assembly and organization, olfactory perception, retinal microglia as well as other microglia types. The neurological and inflammatory manifestations in SjD have been described in depth in literature [70, 71]. For example, the link between SjD and alterations in synaptic signaling within the CNS has been suggested through the Kynurenine Metabolic pathway (KP) interference, which is stimulated by Interferon gamma and interferes with neurotransmission in hippocampus [70]. KP interference of serotonin and glutamate neurotransmitters has been suggested to neurological and psychological symptoms experienced by Sjögren’s patients, such as hyperalgesia and depression [70]. Similarly, dysregulated olfactory perception and olfactory cells have emerged as dysregulated pathways in this analysis. It is worth noting that olfactory dysfunction is a known symptom of SjD, including hyposmia and anosmia [72–74]. This has been attributed to the impact of the autoimmune condition on exocrine glands in nasal passages, leading to dryness, obstruction in nasal passages and reduced olfactory acuity [72–74]. Finally, involvement and dysregulation of glial cells, including retinal and other types of microglia has emerged as a potential player in this study. The chronic inflammation in SjD can trigger the activation of glial cells, especially microglia [75, 76]. In SjD, microglia are known to play a significant role in orchestrating the neuroinflammatory environment, potentially through the KP pathway [70]. The altered neuroinflammatory environment and response by microglia inflammatory factors produced in SjD can disrupt nervous systems normal functioning, leading to some of the common neurological symptoms observed in SjD [71, 77, 78]. The activated microglia can further exacerbate the inflammatory response in the retina, contributing to retinal damage and leading to retinal vasculitis [79–81].

While very little is known and understood about SSA-pathophysiology, some of the genes identified in the SSA-gene signature have been previously implicated in SjD. For example, as reported above, A2M has been previously identified as a potential therapeutic biomarker and a hub gene in Sjögren’s molecular network [60, 61]. Interestingly, two of the lincRNAs identified in our SSA-gene signature, namely, LINC01303 and CORO1A-AS1, have been previously implicated in SjD. For example, LINC01303 has been suggested to play a role in pathophysiology, development, and progression of SjD by modulating expression level of genes involved in interferon and T-cell mediated pathology [82, 83]. On the other hand, CORO1-AS1 has been suggested to contribute to the characteristic inflammation and immune dysfunction observed in SjD [84–86].

Interestingly, we observed three genes were shared between the SSA+ and SSA-gene signatures, namely A2M, DHX58 and ENSG00000250644. These three genes alone were able to achieve an AUC of 0.78, with 73% sensitivity at 82% specificity when distinguishing all samples from controls. Using Sicca only as control, we achieved 72% sensitivity at 82% specificity (Supplementary Fig. 10). A2M, as described before, has been described as a hub gene and implicated in Sjögren’s pathophysiology [60, 61]. DHX58, as described before, is known to play a role in innate immune response and involved in interferon pathways, dysregulation of which has been implicated in SjD [66, 67]. While there is no known relevance of long non-coding RNA ENSG00000250644 in SjD, it may be related to Cathepsin D elevation and inflammation [87]. This may be a potentially novel biomarker that requires further investigation.

When gene expression levels of SSA+ subtype are directly compared with SSA-subtype, pathway analysis of differentially expressed genes point to the upregulation of interferon response pathways in SSA+, while various synaptic and glial pathways were found to be upregulated in SSA-subtype. These results point to the involvement of distinct molecular mechanisms underlying the two Sjögren’s subtypes. Further experiments are needed to corroborate these results.

Lastly, a relatively small sample size and lack of both healthy & patient samples from each collection site are admitted limitations of this study. Yet, this proof-of-concept study highlights the tremendous potential of salivary EVs and its long RNA cargo in biomarker discovery and liquid biopsy. While the biological pathways identified in SSA+ are in strong agreement with known literature, pathways analysis of SSA-samples provide novel insights on the molecular mechanisms underlying this less understood condition. Nevertheless, the biomarkers identified in this pilot study require further experimental exploration and clinical validation in a larger sample cohort, which is currently under investigation.

## Methods

### Sample Collection

Saliva samples were collected and processed at the collection sites using IRB protocol (IRB # 12132). Samples were collected from two sites: participants were recruited from the Research Institute for Systemic Autoimmune Diseases, Athens, Greece and the Division of Oral Medicine, Tufts School of Dental Medicine (Boston MA, USA). Informed consent was obtained from all participants prior to sample collection, and all procedures were conducted in compliance with the ethical standards outlined in the Declaration of Helsinki. Prior to sample collection, patients were advised to abstain from alcohol for 24 hours and refrain from brushing teeth, eating, drinking, or smoking for 1.5 hours. Consumption of water was permitted up to 1 hour before collection.**\**

The exclusion criteria for SjD subjects are outlined as follows: individuals with an active severe infection or chronic infections such as HIV, hepatitis B, hepatitis C, or tuberculosis were excluded. To ensure the biomarker signature is specific for SjD, patients with pre-existing connective tissue or autoimmune diseases like systemic lupus erythematosus (SLE), rheumatoid arthritis, systemic sclerosis, or sarcoidosis were deemed ineligible. Pregnant or breastfeeding females, individuals with a history of head/neck radiation, and those with graft vs. host disease were also excluded from participation.

### Sample processing and Extracellular Vesicle RNA Isolation

Stimulated whole saliva was collected by instructing participants to chew on paraffin wax (Cat no. 3978-289, Benco Dental,) for up to 10 minutes and deposit saliva into a vial treated with RNase inhibitor (Cat no. NC1081844, Fisher Scientific). After samples were collected, they were centrifuged at 2,600 x g for 15 minutes to remove cells. Subsequently, all de-identified samples were stored at −80 °C until further processing. 0.5 mL of patient’s saliva was diluted with 0.5 mL of PBS prior to filtering with 0.8µm syringe filters (catalog no. SLAAR33SB, Fisher Scientific) to exclude extracellular vesicles larger than 800 nm in diameter. Salivary EVs, including exosomes, were isolated, and their total RNAs were extracted by Exosome Diagnostics (Waltham, MA, USA) using the ExoLution™ isolation platform [88]. The RNA yield and size distribution of each sample was then assessed using Agilent’s Bioanalyzer RNA 6000 Pico assay (Catalog no. 5067-1513, Agilent Technologies).

### Whole Transcriptome Library Construction

Exosome Diagnostics (Waltham, MA) conducted the construction and sequencing of Total RNA-Seq libraries using their exclusive EV long RNA-Seq platform. Briefly, EV RNA samples extracted from saliva underwent DNase treatment to eliminate any residual DNA. Following DNA digestion, synthetic RNA spike-in controls (ERCC spike-in mix, catalog no. 4456740, Thermo Fisher Scientific) were introduced into each sample. The EV RNA/ERCC blend was then fragmented for an optimal insert size and reverse transcribed using random priming. Then, second strand synthesis and A-Tailing were performed to facilitate adapter ligation. This was followed by two rounds of purification utilizing AMPureXP® beads (catalog no. A63881, Beckman Coulter). The final libraries were then amplified using a high-fidelity polymerase to a desired concentration for sequencing and purified once more using AMPure-XP® to remove any primer dimers. The Agilent Tape Station High Sensitivity D1000 Screen Tape Assay (catalog no. 5067-5584, Agilent Technologies) and Qubit 1X dsDNA HS Assay Kit (catalog no. Q33231, Thermo Fisher Scientific) were used to quantify libraries. Finally, libraries were normalized and pooled for target enrichment.

### Target Enrichment

Libraries pooled by mass were hybridized to custom human whole exome plus UTR and lncRNAs biotinylated probe set (Agilent Technologies) with a 1-hour hybridization time according to manufacturer recommendations (catalog no. G9987A, Agilent Technologies). Hybridized DNA was then captured using streptavidin-coated beads at 70 °C through a series of wash buffers. PCR amplification was performed to generate enough material for sequencing and then purified using a single round of AMPureXP® beads (catalog no. A63881, Beckman Coulter). Libraries were quantified using the Agilent Tape station High Sensitivity D1000 Screen Tape Assay (catalog no. 5067-5584, Agilent Technologies) and Qubit 1X dsDNA HS Assay Kit (catalog no. Q33231, Thermo Fisher Scientific). Final captures were diluted to a desired concentration and sequenced on Illumina® NextSeq500 or NextSeq2000 sequencers using 2 × 75 cycles read length chemistry.

### Bioinformatics

Sequencing reads (2 × 75 bp, paired end) were aligned to the human reference genome (GRCh38) using STAR (v2.7.9a), guided by Ensembl GTF release 106 annotations [43]. The resulting alignments were processed with Salmon (v1.9.0) to quantify transcript abundance, incorporating both sequence-specific and GC bias corrections to compute transcript-level read counts and transcripts per million (TPM) values [89]. Transcript-level quantifications were aggregated to the gene level using the tximport package in Bioconductor. DESeq2 was then used to perform differential gene expression (DEx) analysis on raw read counts [90]. P-values were adjusted for multiple testing using the Benjamini-Hochberg method, and genes were considered statistically significant if they had an adjusted p-value < 0.05 and a |log2 fold change| > 1.

To explore biological relevance, pathway enrichment analysis was performed using the clusterProfiler package with gene sets from the MSigDB database [44, 45]. Additionally, tissue deconvolution was conducted using the DeconRNASeq package, referencing expression profiles from GTEx and the Tabula Sapiens v1.0 (TSP) database [91, 92].

To control potential technical artifacts arising from differences in sample collection sites, we used DESeq2 to identify genes that varied significantly between sites within the same disease group. These site-specific genes were excluded from downstream biomarker discovery to reduce false positives.

We implemented a 10-repeat stratified 5-fold nested cross-validation (CV) framework to identify robust biomarkers and prevent data leakage. Data were stratified by condition, and each outer CV fold contained a complete inner CV loop used for feature selection and model training.

Within each inner CV, feature selection was performed using a multi-step pipeline, including variance filtering, false positive rate (FPR) filtering, Lasso regression, and Boruta selection. A grid search was used to tune model hyperparameters and constrain the final feature set to fewer than 10 predictors.

The best-performing model from the inner CV was retrained on the full training set of the outer CV and then evaluated on the corresponding test fold. This process was repeated 50 times (10 repeats × 5 folds), and predictions from all outer test folds were aggregated to compute the area under the receiver operating characteristic curve (AUC-ROC), which was used to assess classifier performance and generalizability.

## Code Availability

https://github.com/exosomedx/sjogren_biomarker_discovery.git

