## Supplemental Figures and Tables for "Salivary Extracellular Vesicle RNA Profiling Reveals Biomarkers for Sjögren’s"

### Slide 1
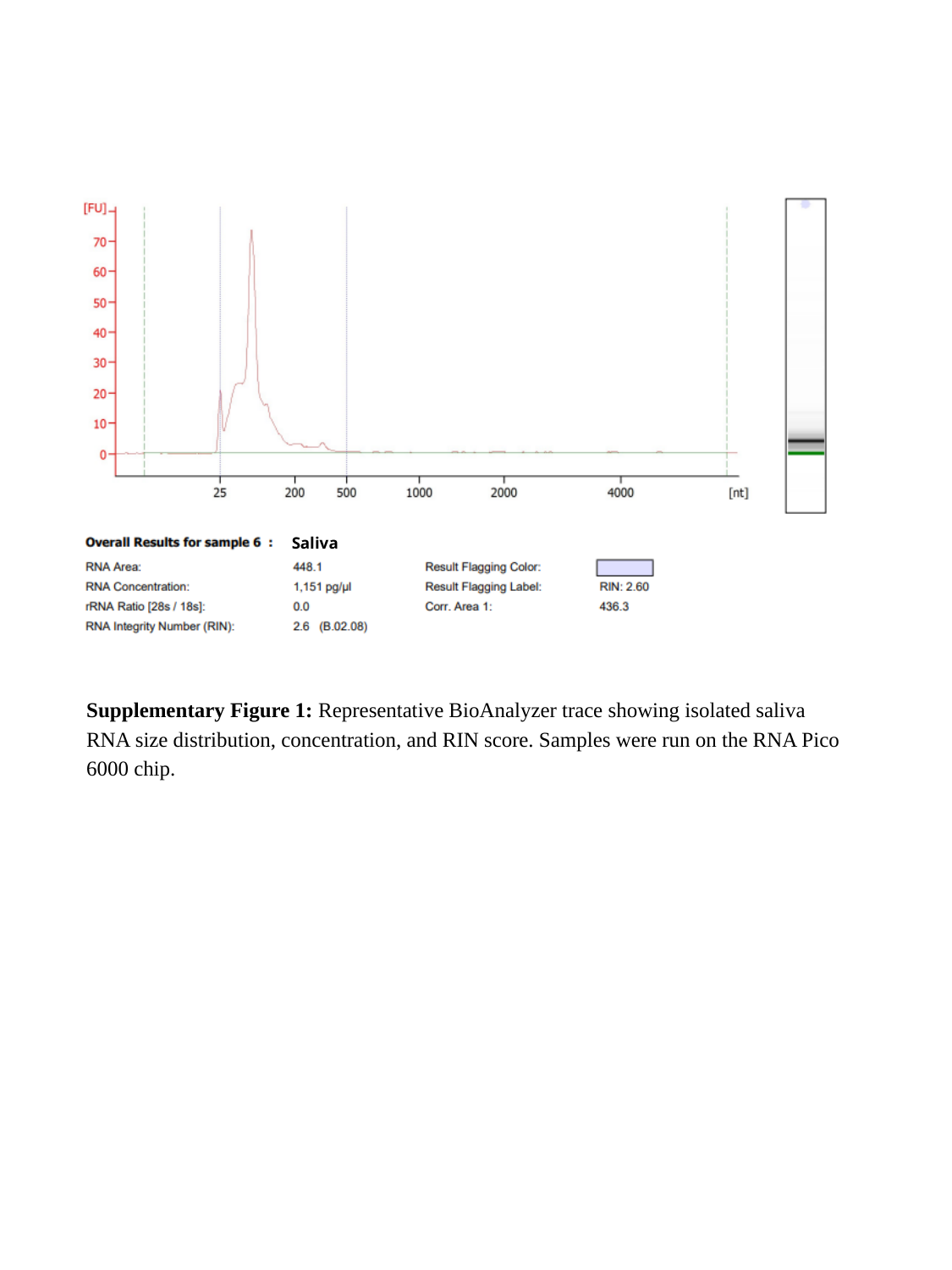

Saliva
Supplementary Figure 1: Representative BioAnalyzer trace showing isolated saliva RNA size distribution, concentration, and RIN score. Samples were run on the RNA Pico 6000 chip.

### Slide 2
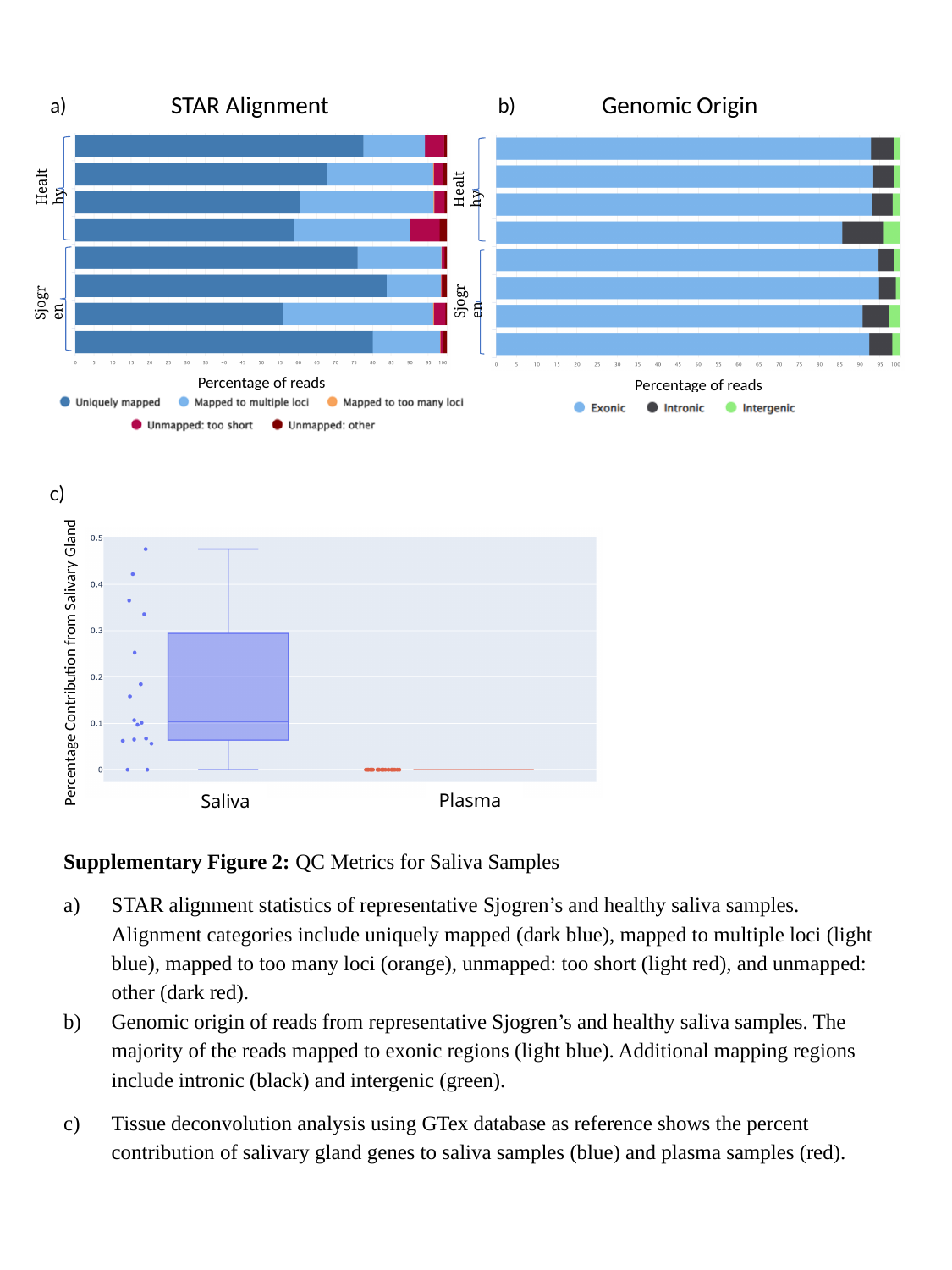

STAR Alignment
Genomic Origin
a)
b)
Healthy
Healthy
Sjogren
Sjogren
Percentage of reads
Percentage of reads
c)
Percentage Contribution from Salivary Gland
Plasma
Saliva
Supplementary Figure 2: QC Metrics for Saliva Samples
STAR alignment statistics of representative Sjogren’s and healthy saliva samples. Alignment categories include uniquely mapped (dark blue), mapped to multiple loci (light blue), mapped to too many loci (orange), unmapped: too short (light red), and unmapped: other (dark red).
Genomic origin of reads from representative Sjogren’s and healthy saliva samples. The majority of the reads mapped to exonic regions (light blue). Additional mapping regions include intronic (black) and intergenic (green).
Tissue deconvolution analysis using GTex database as reference shows the percent contribution of salivary gland genes to saliva samples (blue) and plasma samples (red).

### Slide 3
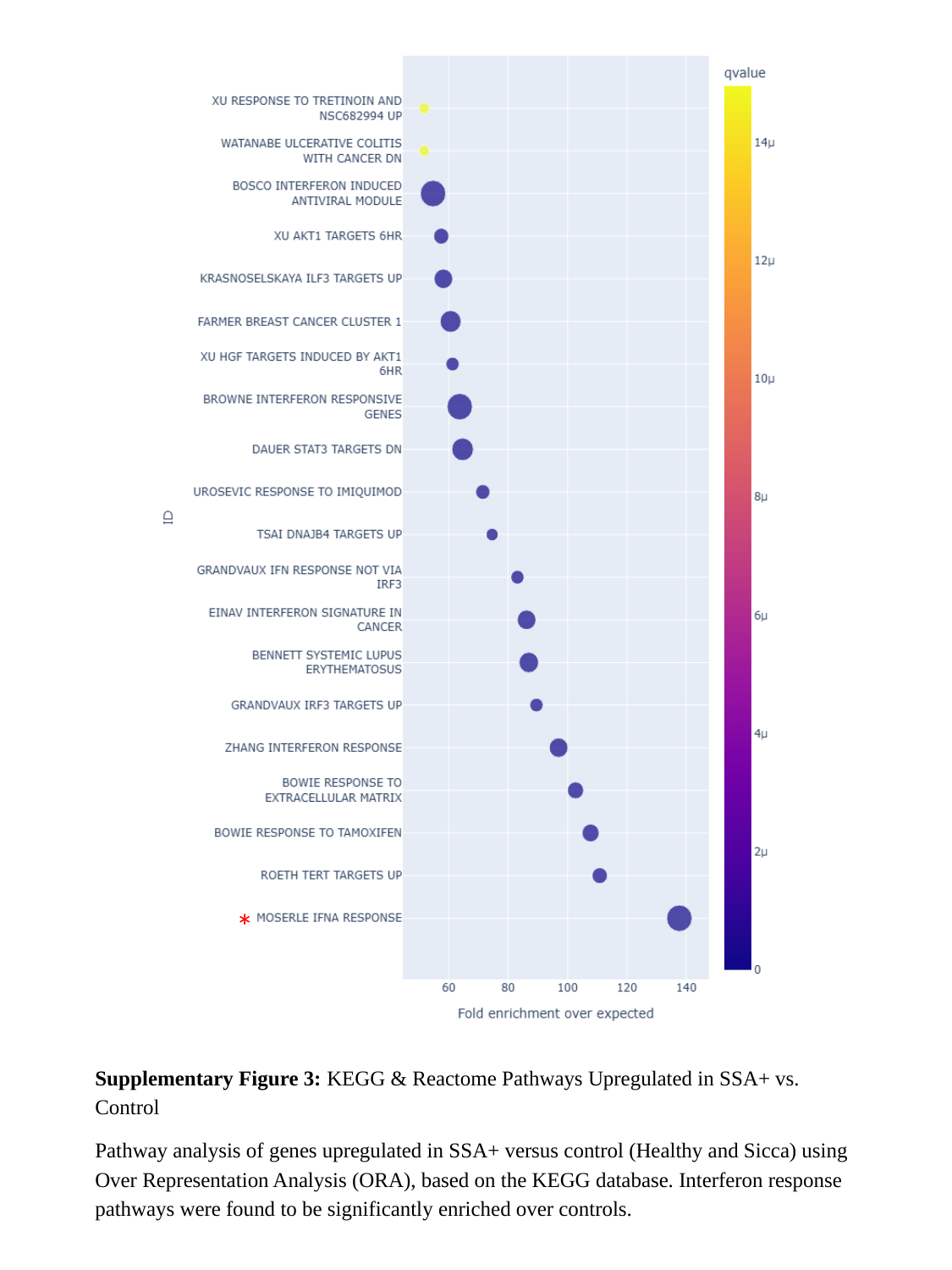

*
Supplementary Figure 3: KEGG & Reactome Pathways Upregulated in SSA+ vs. Control
Pathway analysis of genes upregulated in SSA+ versus control (Healthy and Sicca) using Over Representation Analysis (ORA), based on the KEGG database. Interferon response pathways were found to be significantly enriched over controls.

### Slide 4
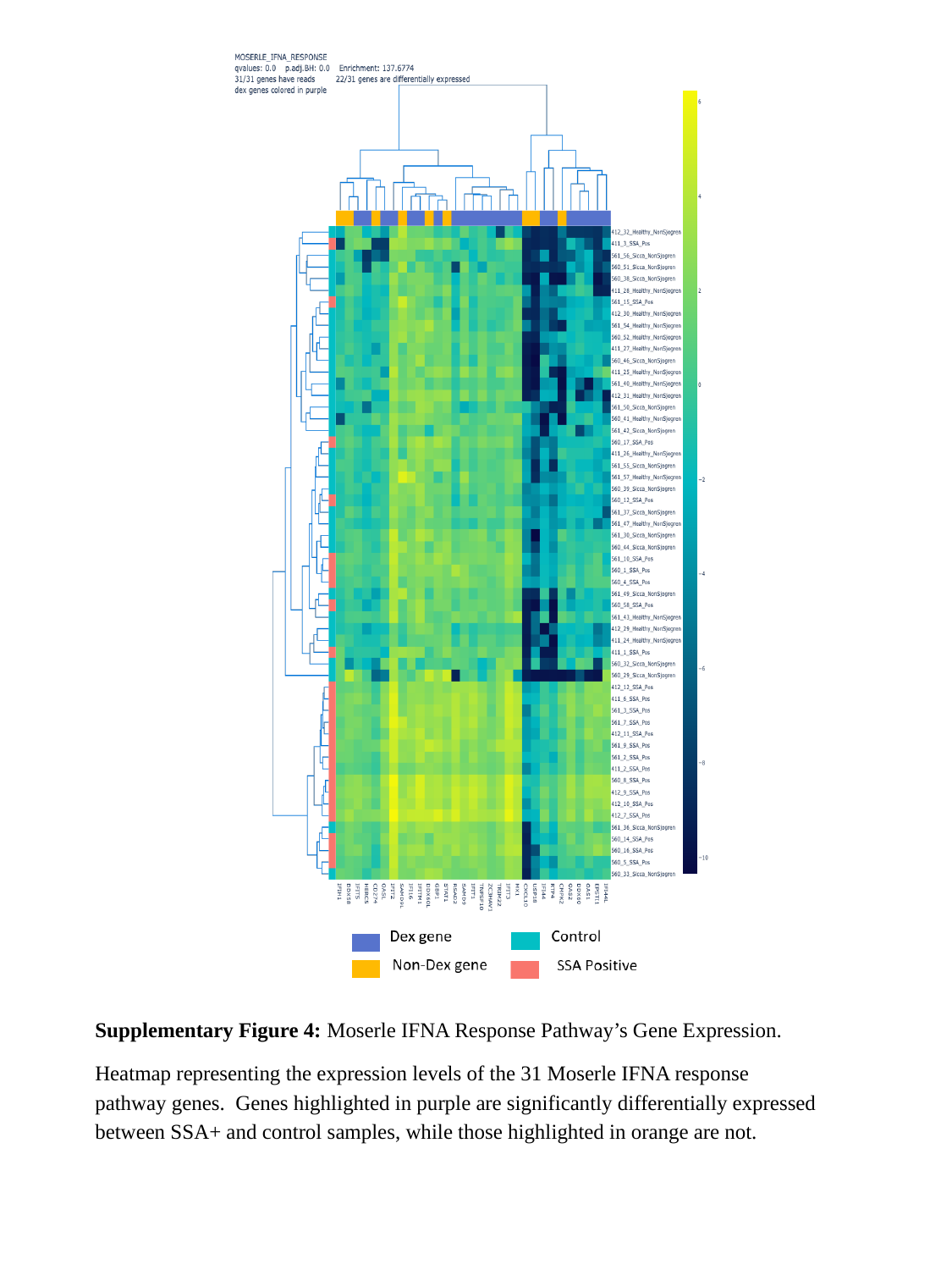

Supplementary Figure 4: Moserle IFNA Response Pathway’s Gene Expression.
Heatmap representing the expression levels of the 31 Moserle IFNA response pathway genes. Genes highlighted in purple are significantly differentially expressed between SSA+ and control samples, while those highlighted in orange are not.

### Slide 5
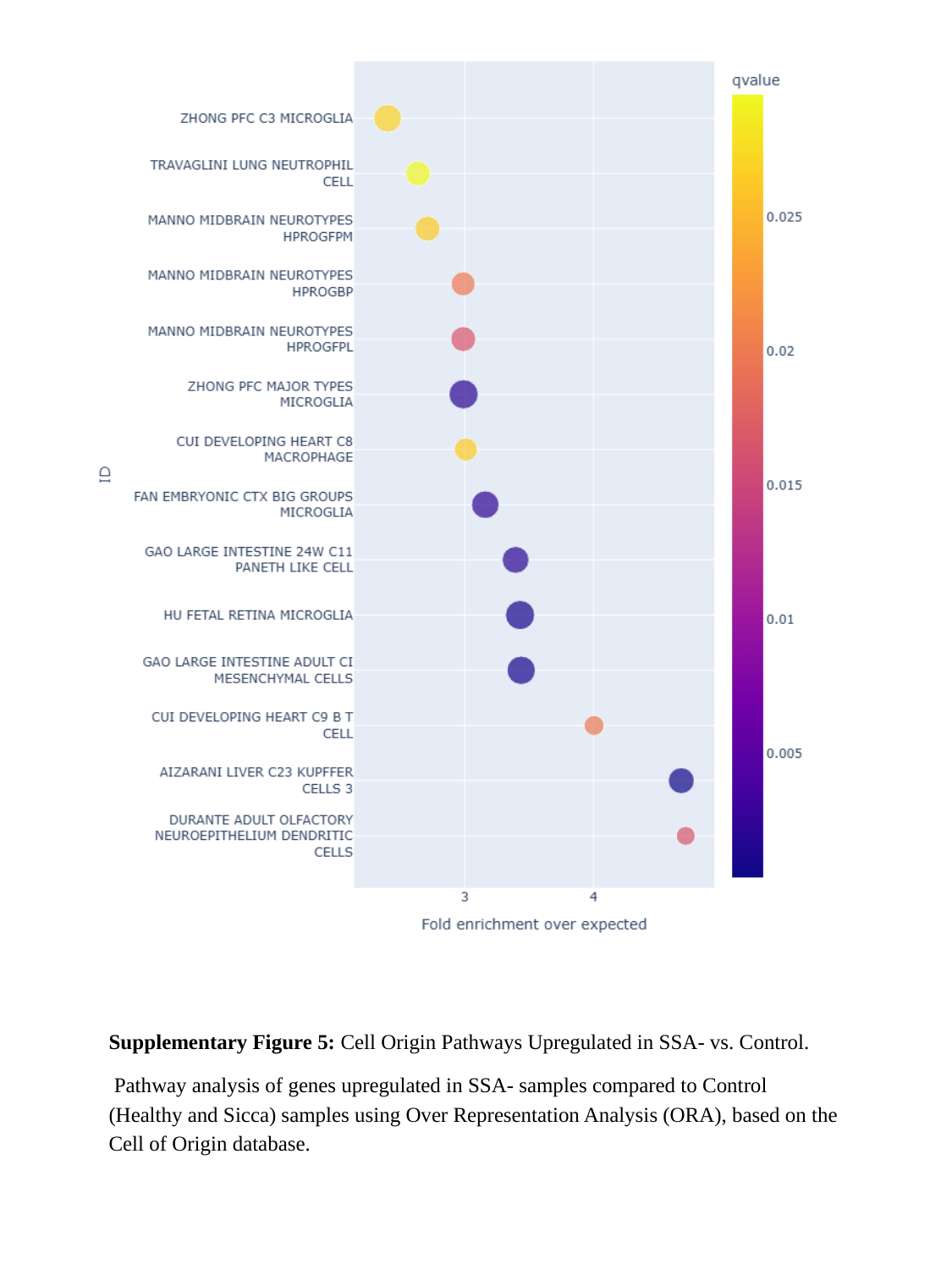

Supplementary Figure 5: Cell Origin Pathways Upregulated in SSA- vs. Control.
 Pathway analysis of genes upregulated in SSA- samples compared to Control (Healthy and Sicca) samples using Over Representation Analysis (ORA), based on the Cell of Origin database.

### Slide 6
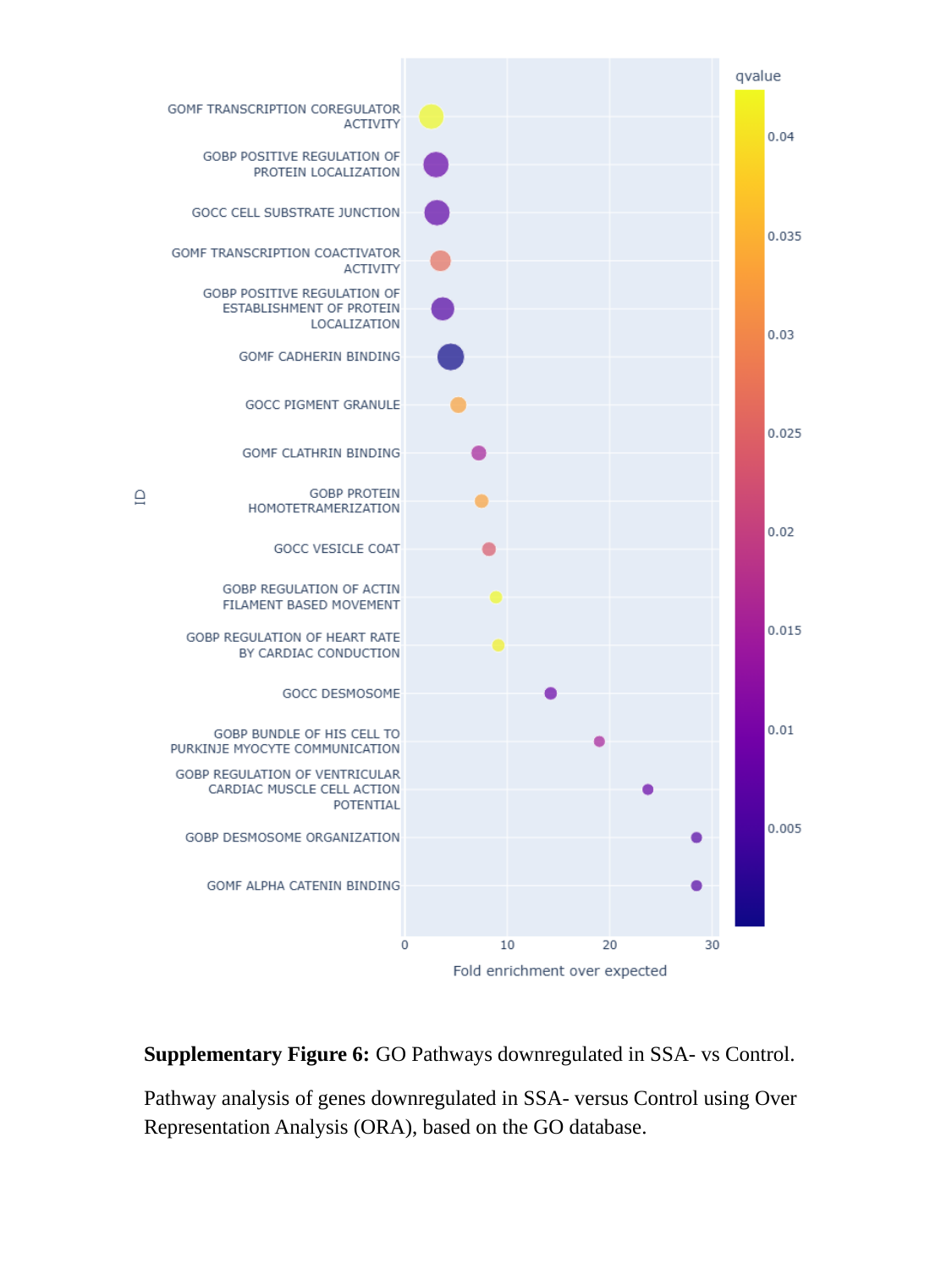

Supplementary Figure 6: GO Pathways downregulated in SSA- vs Control.
Pathway analysis of genes downregulated in SSA- versus Control using Over Representation Analysis (ORA), based on the GO database.

### Slide 7
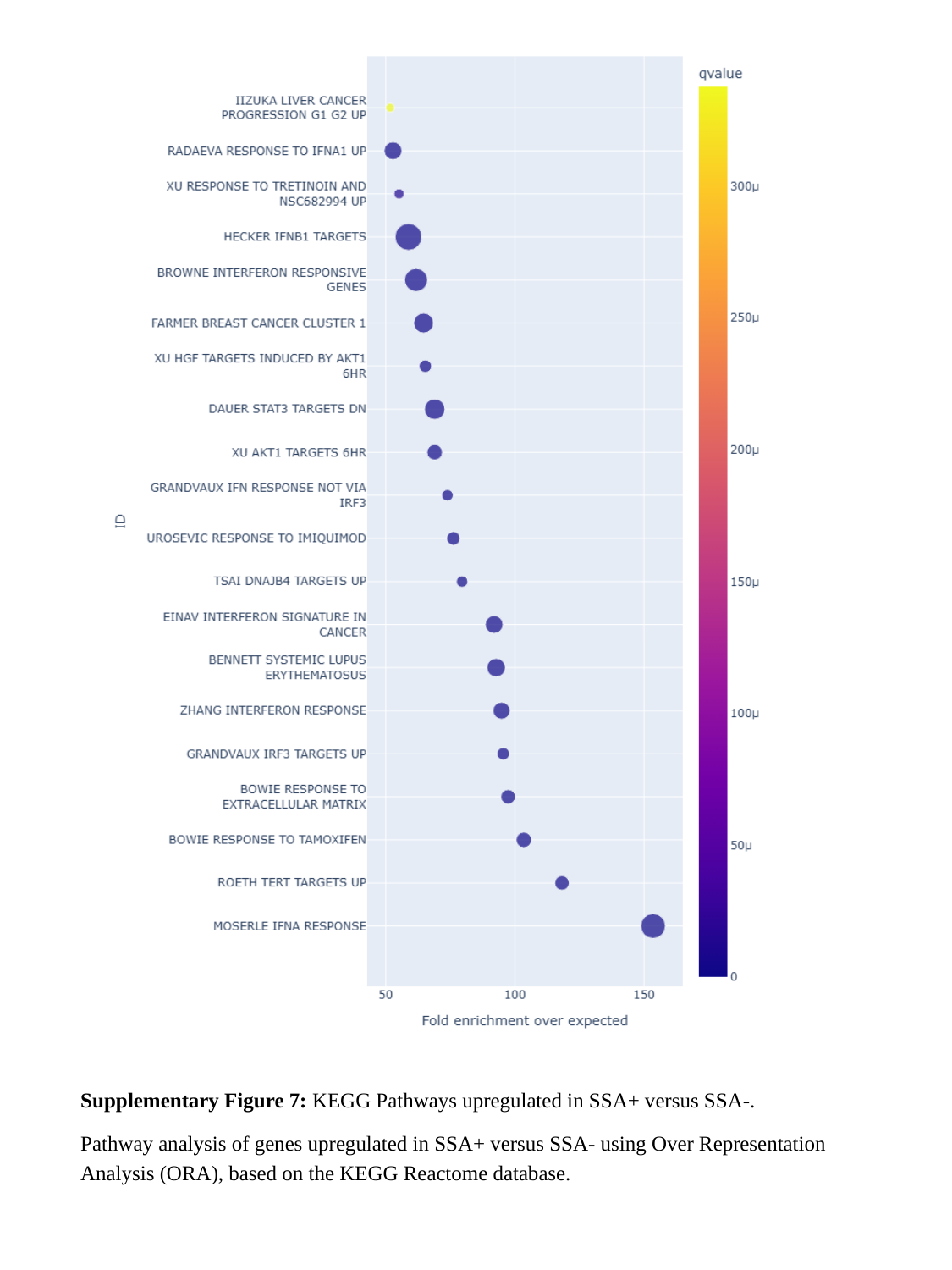

Supplementary Figure 7: KEGG Pathways upregulated in SSA+ versus SSA-.
Pathway analysis of genes upregulated in SSA+ versus SSA- using Over Representation Analysis (ORA), based on the KEGG Reactome database.

### Slide 8
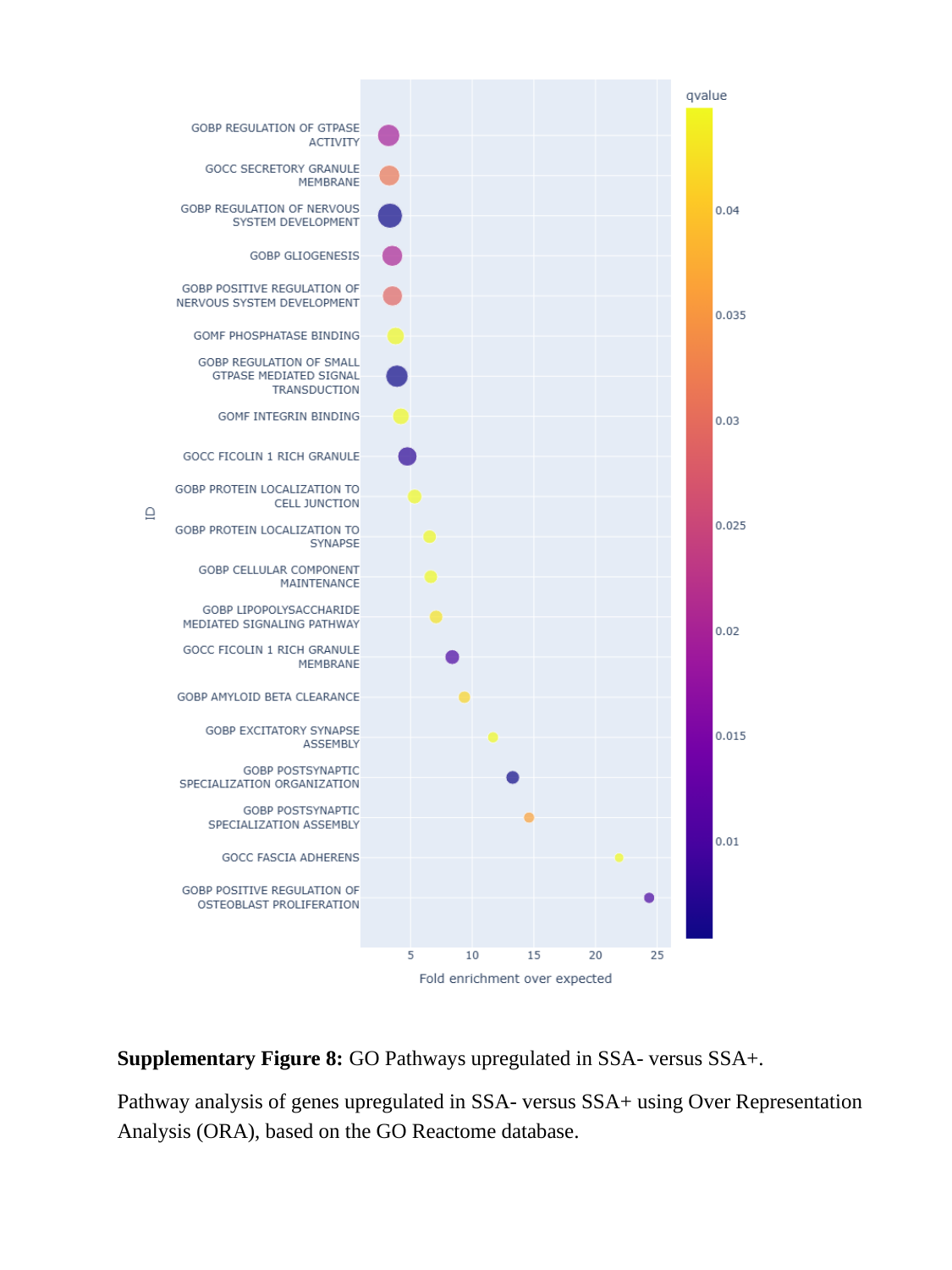

Supplementary Figure 8: GO Pathways upregulated in SSA- versus SSA+.
Pathway analysis of genes upregulated in SSA- versus SSA+ using Over Representation Analysis (ORA), based on the GO Reactome database.

### Slide 9
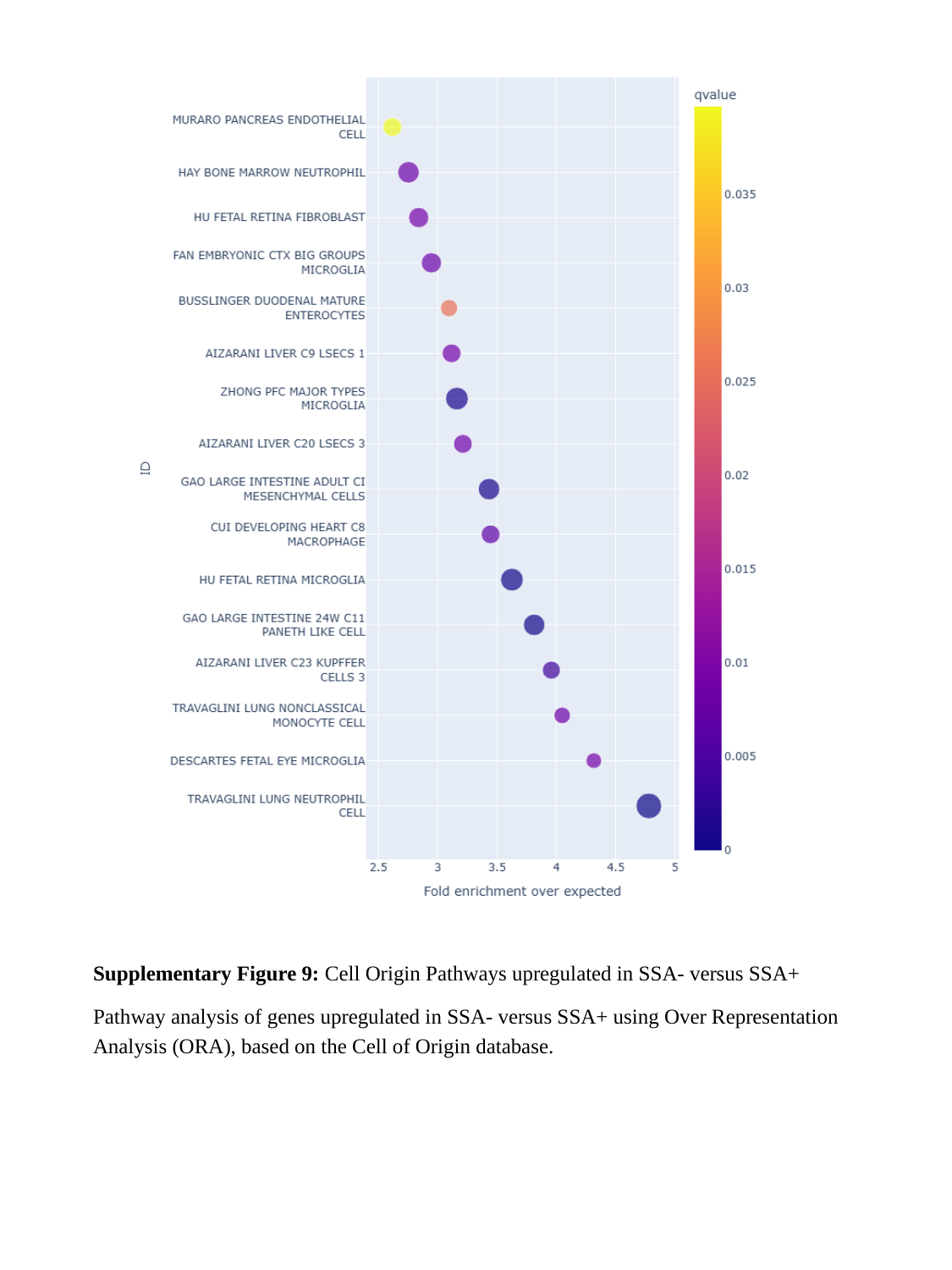

Supplementary Figure 9: Cell Origin Pathways upregulated in SSA- versus SSA+
Pathway analysis of genes upregulated in SSA- versus SSA+ using Over Representation Analysis (ORA), based on the Cell of Origin database.

### Slide 10
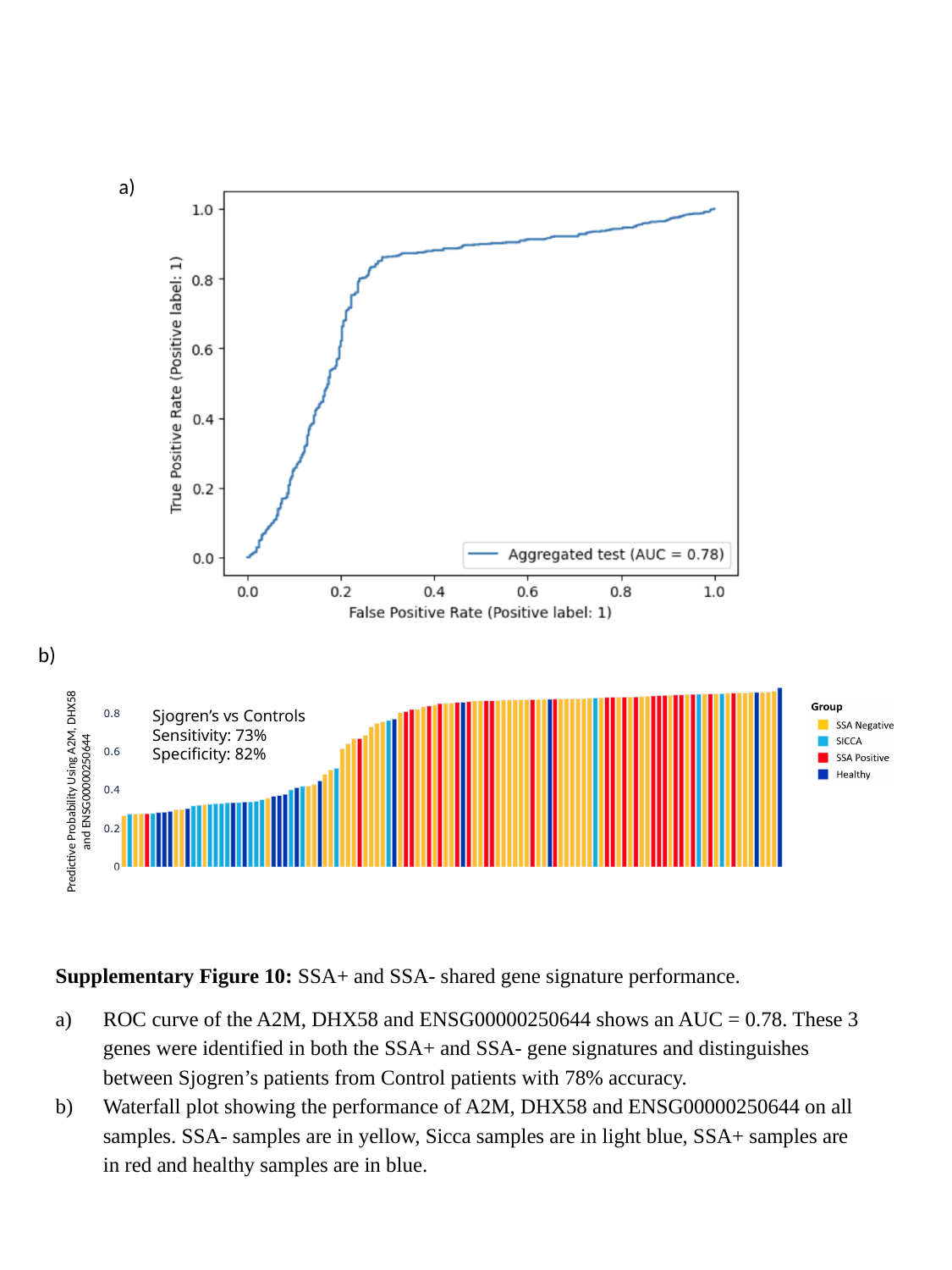

a)
b)
Sjogren’s vs Controls
Sensitivity: 73%
Specificity: 82%
Predictive Probability Using A2M, DHX58 and ENSG00000250644
Supplementary Figure 10: SSA+ and SSA- shared gene signature performance.
ROC curve of the A2M, DHX58 and ENSG00000250644 shows an AUC = 0.78. These 3 genes were identified in both the SSA+ and SSA- gene signatures and distinguishes between Sjogren’s patients from Control patients with 78% accuracy.
Waterfall plot showing the performance of A2M, DHX58 and ENSG00000250644 on all samples. SSA- samples are in yellow, Sicca samples are in light blue, SSA+ samples are in red and healthy samples are in blue.

### Slide 11
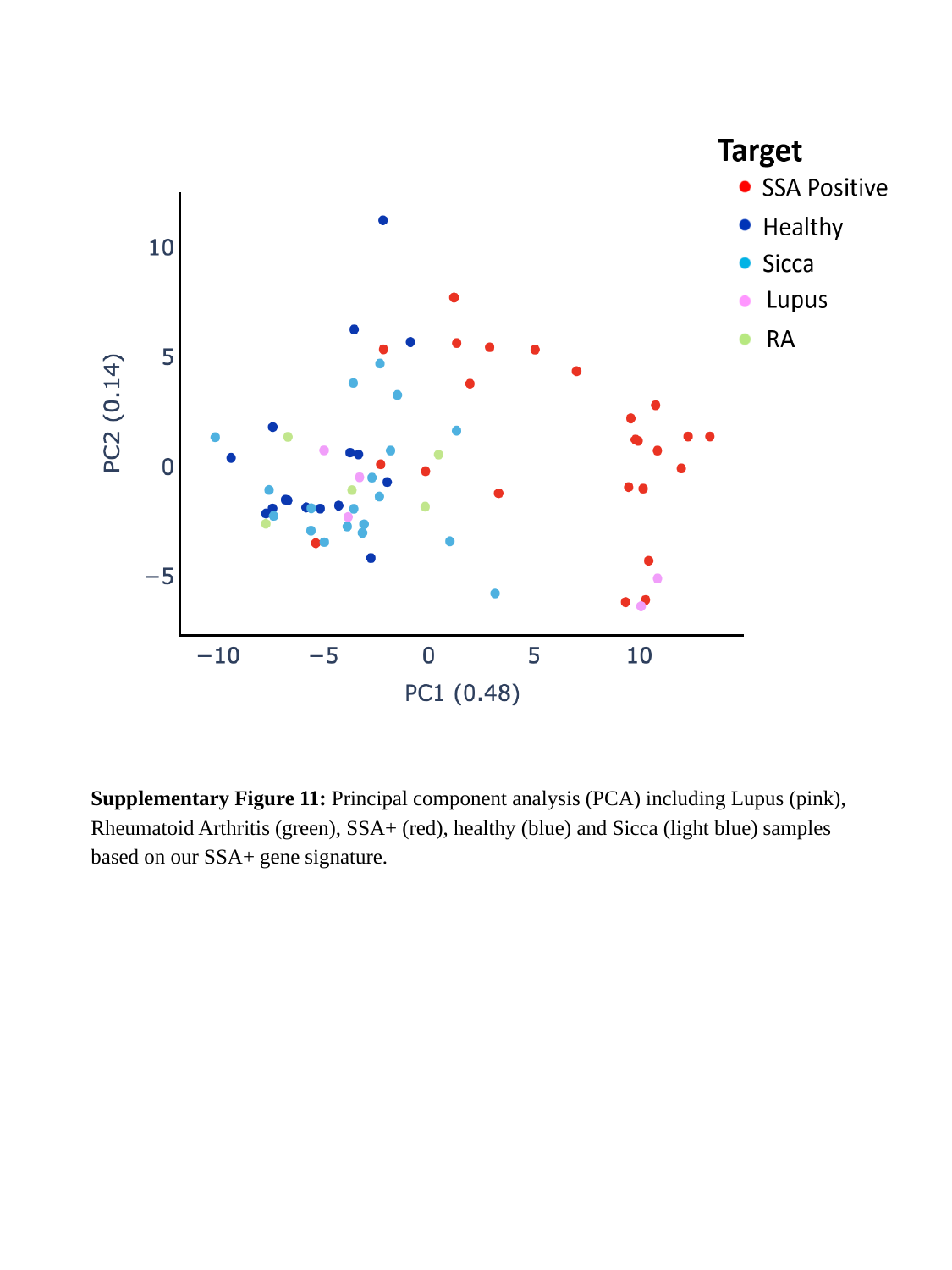

Supplementary Figure 11: Principal component analysis (PCA) including Lupus (pink), Rheumatoid Arthritis (green), SSA+ (red), healthy (blue) and Sicca (light blue) samples based on our SSA+ gene signature.

### Slide 12
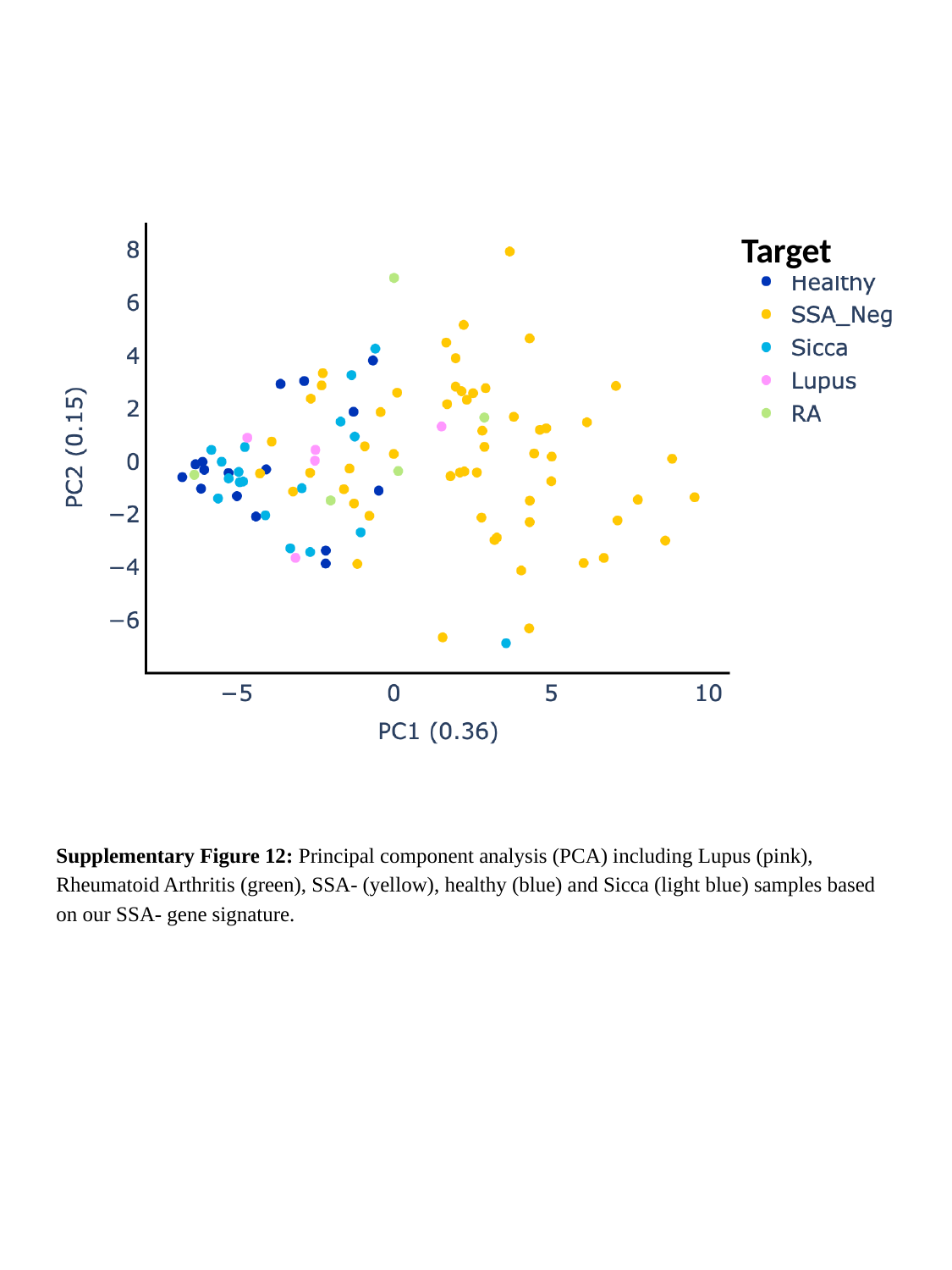

Target
Supplementary Figure 12: Principal component analysis (PCA) including Lupus (pink), Rheumatoid Arthritis (green), SSA- (yellow), healthy (blue) and Sicca (light blue) samples based on our SSA- gene signature.

### Slide 13
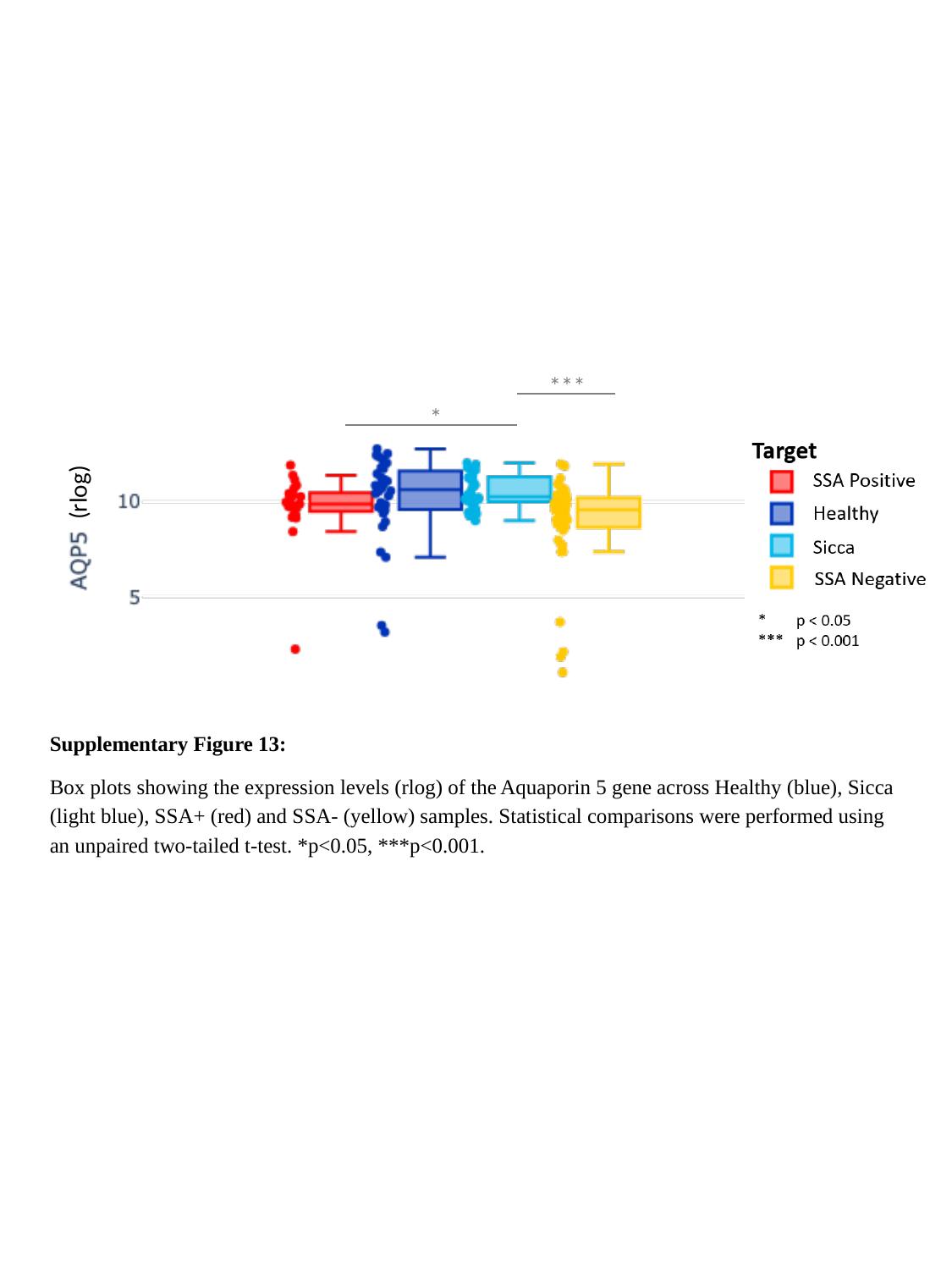

***
*
(rlog)
Supplementary Figure 13:
Box plots showing the expression levels (rlog) of the Aquaporin 5 gene across Healthy (blue), Sicca (light blue), SSA+ (red) and SSA- (yellow) samples. Statistical comparisons were performed using an unpaired two-tailed t-test. *p<0.05, ***p<0.001.

### Slide 14
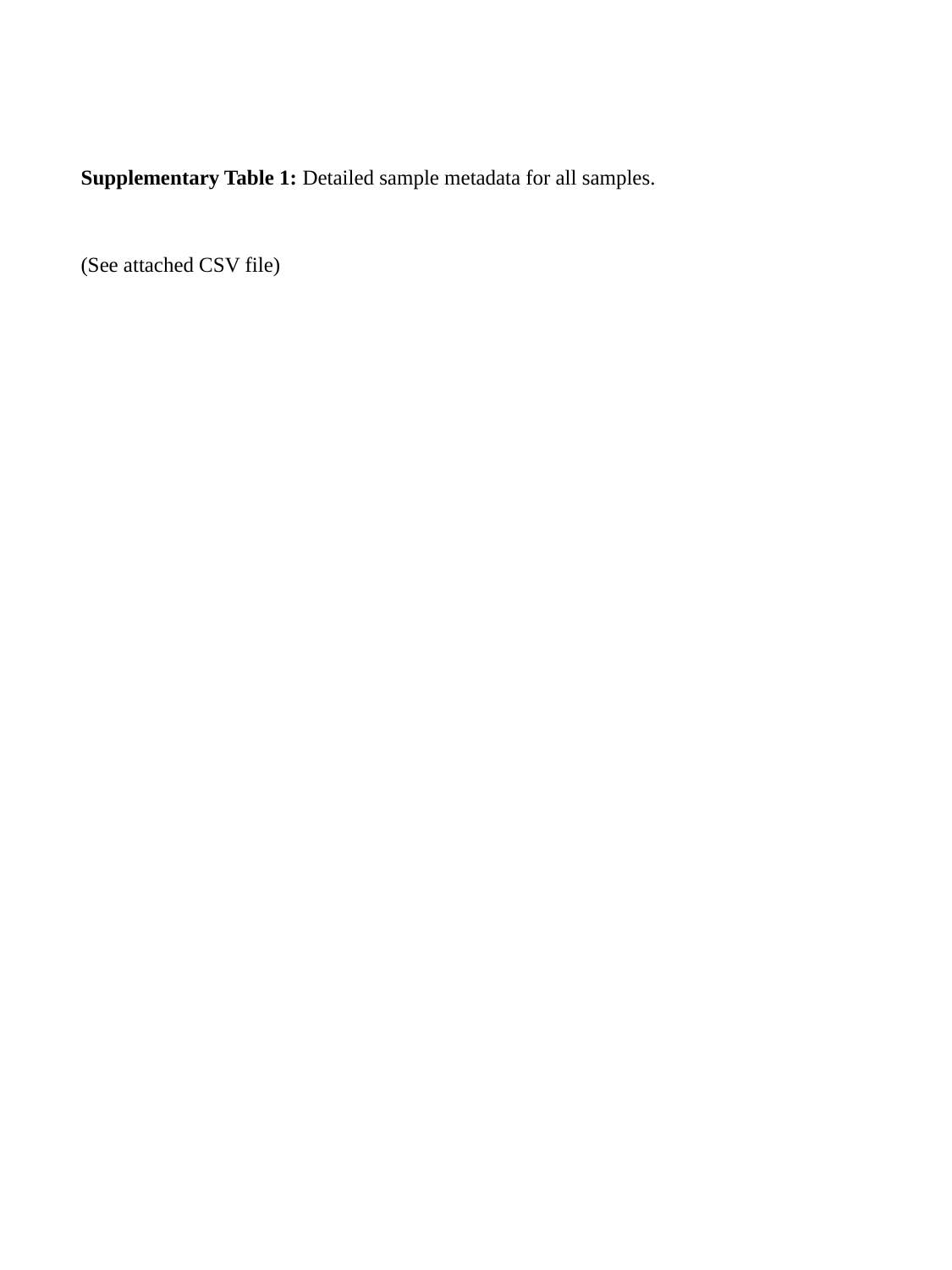

Supplementary Table 1: Detailed sample metadata for all samples.
(See attached CSV file)

### Slide 15
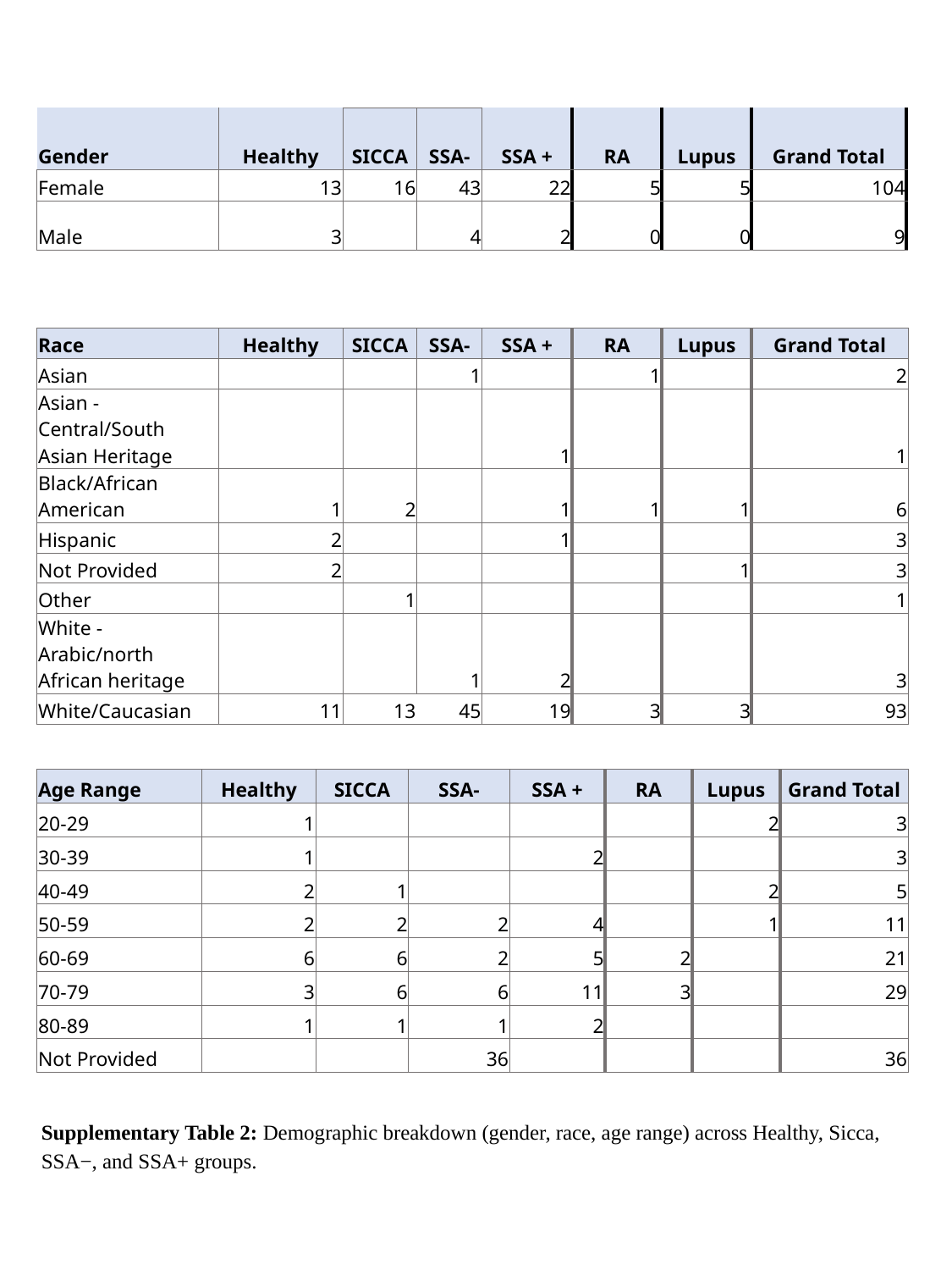

| Gender | Healthy | SICCA | SSA- | SSA + | RA | Lupus | Grand Total |
| --- | --- | --- | --- | --- | --- | --- | --- |
| Female | 13 | 16 | 43 | 22 | 5 | 5 | 104 |
| Male | 3 | | 4 | 2 | 0 | 0 | 9 |
| Race | Healthy | SICCA | SSA- | SSA + | RA | Lupus | Grand Total |
| --- | --- | --- | --- | --- | --- | --- | --- |
| Asian | | | 1 | | 1 | | 2 |
| Asian - Central/South Asian Heritage | | | | 1 | | | 1 |
| Black/African American | 1 | 2 | | 1 | 1 | 1 | 6 |
| Hispanic | 2 | | | 1 | | | 3 |
| Not Provided | 2 | | | | | 1 | 3 |
| Other | | 1 | | | | | 1 |
| White - Arabic/north African heritage | | | 1 | 2 | | | 3 |
| White/Caucasian | 11 | 13 | 45 | 19 | 3 | 3 | 93 |
| Age Range | Healthy | SICCA | SSA- | SSA + | RA | Lupus | Grand Total |
| --- | --- | --- | --- | --- | --- | --- | --- |
| 20-29 | 1 | | | | | 2 | 3 |
| 30-39 | 1 | | | 2 | | | 3 |
| 40-49 | 2 | 1 | | | | 2 | 5 |
| 50-59 | 2 | 2 | 2 | 4 | | 1 | 11 |
| 60-69 | 6 | 6 | 2 | 5 | 2 | | 21 |
| 70-79 | 3 | 6 | 6 | 11 | 3 | | 29 |
| 80-89 | 1 | 1 | 1 | 2 | | | |
| Not Provided | | | 36 | | | | 36 |
Supplementary Table 2: Demographic breakdown (gender, race, age range) across Healthy, Sicca, SSA−, and SSA+ groups.

### Slide 16
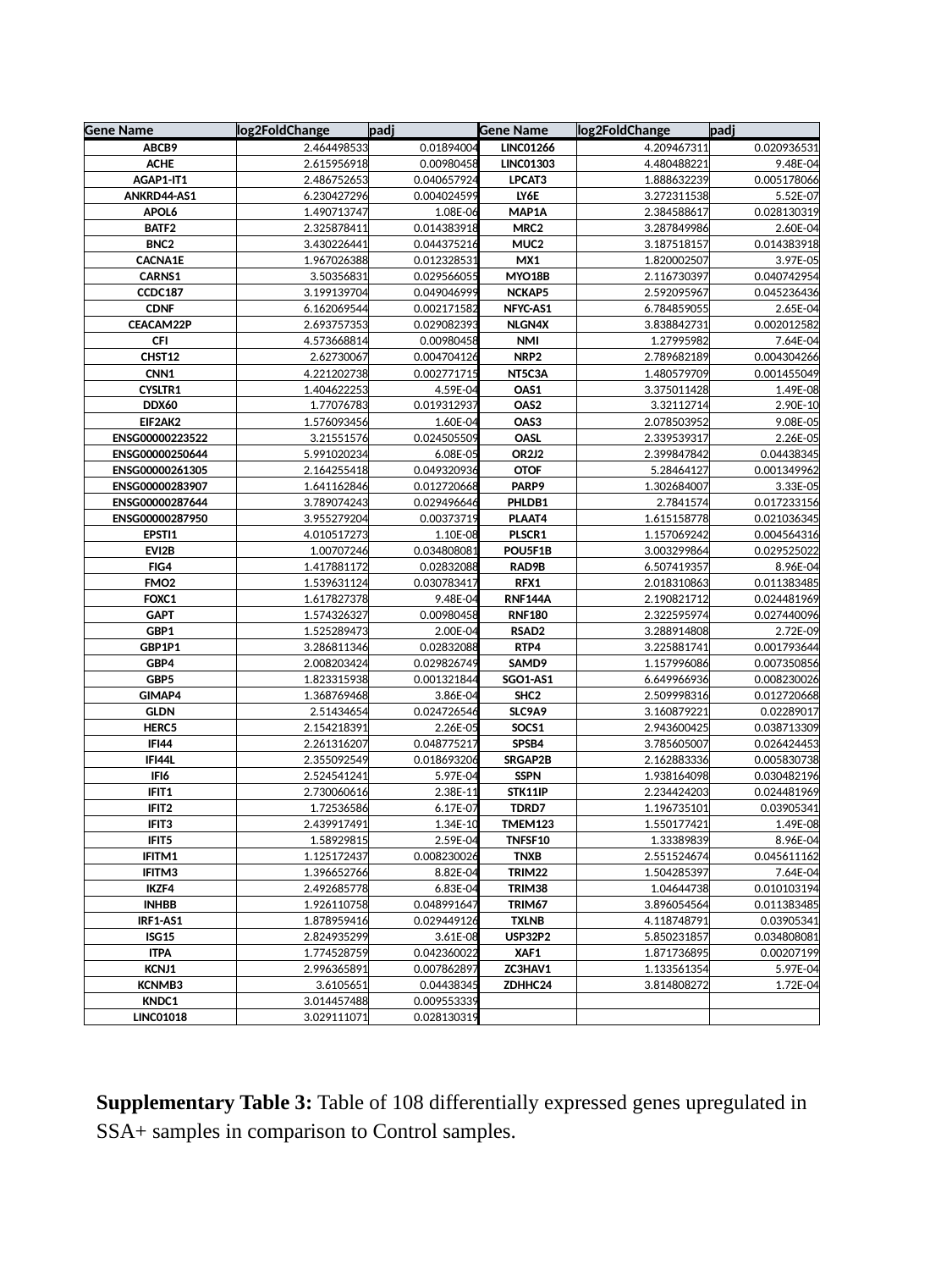

| Gene Name | log2FoldChange | padj | Gene Name | log2FoldChange | padj |
| --- | --- | --- | --- | --- | --- |
| ABCB9 | 2.464498533 | 0.01894004 | LINC01266 | 4.209467311 | 0.020936531 |
| ACHE | 2.615956918 | 0.00980458 | LINC01303 | 4.480488221 | 9.48E-04 |
| AGAP1-IT1 | 2.486752653 | 0.040657924 | LPCAT3 | 1.888632239 | 0.005178066 |
| ANKRD44-AS1 | 6.230427296 | 0.004024599 | LY6E | 3.272311538 | 5.52E-07 |
| APOL6 | 1.490713747 | 1.08E-06 | MAP1A | 2.384588617 | 0.028130319 |
| BATF2 | 2.325878411 | 0.014383918 | MRC2 | 3.287849986 | 2.60E-04 |
| BNC2 | 3.430226441 | 0.044375216 | MUC2 | 3.187518157 | 0.014383918 |
| CACNA1E | 1.967026388 | 0.012328531 | MX1 | 1.820002507 | 3.97E-05 |
| CARNS1 | 3.50356831 | 0.029566055 | MYO18B | 2.116730397 | 0.040742954 |
| CCDC187 | 3.199139704 | 0.049046999 | NCKAP5 | 2.592095967 | 0.045236436 |
| CDNF | 6.162069544 | 0.002171582 | NFYC-AS1 | 6.784859055 | 2.65E-04 |
| CEACAM22P | 2.693757353 | 0.029082393 | NLGN4X | 3.838842731 | 0.002012582 |
| CFI | 4.573668814 | 0.00980458 | NMI | 1.27995982 | 7.64E-04 |
| CHST12 | 2.62730067 | 0.004704126 | NRP2 | 2.789682189 | 0.004304266 |
| CNN1 | 4.221202738 | 0.002771715 | NT5C3A | 1.480579709 | 0.001455049 |
| CYSLTR1 | 1.404622253 | 4.59E-04 | OAS1 | 3.375011428 | 1.49E-08 |
| DDX60 | 1.77076783 | 0.019312937 | OAS2 | 3.32112714 | 2.90E-10 |
| EIF2AK2 | 1.576093456 | 1.60E-04 | OAS3 | 2.078503952 | 9.08E-05 |
| ENSG00000223522 | 3.21551576 | 0.024505509 | OASL | 2.339539317 | 2.26E-05 |
| ENSG00000250644 | 5.991020234 | 6.08E-05 | OR2J2 | 2.399847842 | 0.04438345 |
| ENSG00000261305 | 2.164255418 | 0.049320936 | OTOF | 5.28464127 | 0.001349962 |
| ENSG00000283907 | 1.641162846 | 0.012720668 | PARP9 | 1.302684007 | 3.33E-05 |
| ENSG00000287644 | 3.789074243 | 0.029496646 | PHLDB1 | 2.7841574 | 0.017233156 |
| ENSG00000287950 | 3.955279204 | 0.00373719 | PLAAT4 | 1.615158778 | 0.021036345 |
| EPSTI1 | 4.010517273 | 1.10E-08 | PLSCR1 | 1.157069242 | 0.004564316 |
| EVI2B | 1.00707246 | 0.034808081 | POU5F1B | 3.003299864 | 0.029525022 |
| FIG4 | 1.417881172 | 0.02832088 | RAD9B | 6.507419357 | 8.96E-04 |
| FMO2 | 1.539631124 | 0.030783417 | RFX1 | 2.018310863 | 0.011383485 |
| FOXC1 | 1.617827378 | 9.48E-04 | RNF144A | 2.190821712 | 0.024481969 |
| GAPT | 1.574326327 | 0.00980458 | RNF180 | 2.322595974 | 0.027440096 |
| GBP1 | 1.525289473 | 2.00E-04 | RSAD2 | 3.288914808 | 2.72E-09 |
| GBP1P1 | 3.286811346 | 0.02832088 | RTP4 | 3.225881741 | 0.001793644 |
| GBP4 | 2.008203424 | 0.029826749 | SAMD9 | 1.157996086 | 0.007350856 |
| GBP5 | 1.823315938 | 0.001321844 | SGO1-AS1 | 6.649966936 | 0.008230026 |
| GIMAP4 | 1.368769468 | 3.86E-04 | SHC2 | 2.509998316 | 0.012720668 |
| GLDN | 2.51434654 | 0.024726546 | SLC9A9 | 3.160879221 | 0.02289017 |
| HERC5 | 2.154218391 | 2.26E-05 | SOCS1 | 2.943600425 | 0.038713309 |
| IFI44 | 2.261316207 | 0.048775217 | SPSB4 | 3.785605007 | 0.026424453 |
| IFI44L | 2.355092549 | 0.018693206 | SRGAP2B | 2.162883336 | 0.005830738 |
| IFI6 | 2.524541241 | 5.97E-04 | SSPN | 1.938164098 | 0.030482196 |
| IFIT1 | 2.730060616 | 2.38E-11 | STK11IP | 2.234424203 | 0.024481969 |
| IFIT2 | 1.72536586 | 6.17E-07 | TDRD7 | 1.196735101 | 0.03905341 |
| IFIT3 | 2.439917491 | 1.34E-10 | TMEM123 | 1.550177421 | 1.49E-08 |
| IFIT5 | 1.58929815 | 2.59E-04 | TNFSF10 | 1.33389839 | 8.96E-04 |
| IFITM1 | 1.125172437 | 0.008230026 | TNXB | 2.551524674 | 0.045611162 |
| IFITM3 | 1.396652766 | 8.82E-04 | TRIM22 | 1.504285397 | 7.64E-04 |
| IKZF4 | 2.492685778 | 6.83E-04 | TRIM38 | 1.04644738 | 0.010103194 |
| INHBB | 1.926110758 | 0.048991647 | TRIM67 | 3.896054564 | 0.011383485 |
| IRF1-AS1 | 1.878959416 | 0.029449126 | TXLNB | 4.118748791 | 0.03905341 |
| ISG15 | 2.824935299 | 3.61E-08 | USP32P2 | 5.850231857 | 0.034808081 |
| ITPA | 1.774528759 | 0.042360022 | XAF1 | 1.871736895 | 0.00207199 |
| KCNJ1 | 2.996365891 | 0.007862897 | ZC3HAV1 | 1.133561354 | 5.97E-04 |
| KCNMB3 | 3.6105651 | 0.04438345 | ZDHHC24 | 3.814808272 | 1.72E-04 |
| KNDC1 | 3.014457488 | 0.009553339 | | | |
| LINC01018 | 3.029111071 | 0.028130319 | | | |
Supplementary Table 3: Table of 108 differentially expressed genes upregulated in SSA+ samples in comparison to Control samples.

### Slide 17
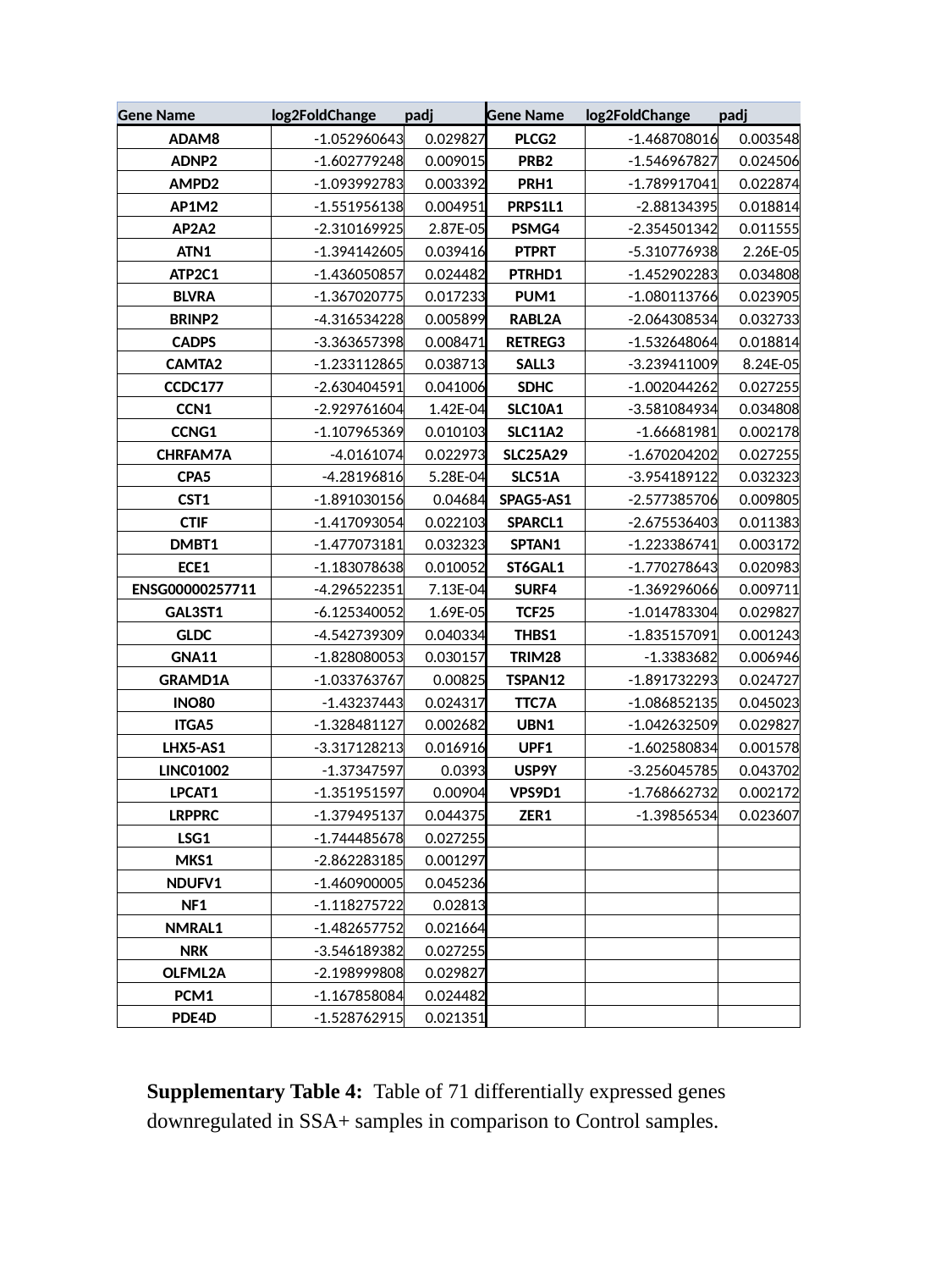

| Gene Name | log2FoldChange | padj | Gene Name | log2FoldChange | padj |
| --- | --- | --- | --- | --- | --- |
| ADAM8 | -1.052960643 | 0.029827 | PLCG2 | -1.468708016 | 0.003548 |
| ADNP2 | -1.602779248 | 0.009015 | PRB2 | -1.546967827 | 0.024506 |
| AMPD2 | -1.093992783 | 0.003392 | PRH1 | -1.789917041 | 0.022874 |
| AP1M2 | -1.551956138 | 0.004951 | PRPS1L1 | -2.88134395 | 0.018814 |
| AP2A2 | -2.310169925 | 2.87E-05 | PSMG4 | -2.354501342 | 0.011555 |
| ATN1 | -1.394142605 | 0.039416 | PTPRT | -5.310776938 | 2.26E-05 |
| ATP2C1 | -1.436050857 | 0.024482 | PTRHD1 | -1.452902283 | 0.034808 |
| BLVRA | -1.367020775 | 0.017233 | PUM1 | -1.080113766 | 0.023905 |
| BRINP2 | -4.316534228 | 0.005899 | RABL2A | -2.064308534 | 0.032733 |
| CADPS | -3.363657398 | 0.008471 | RETREG3 | -1.532648064 | 0.018814 |
| CAMTA2 | -1.233112865 | 0.038713 | SALL3 | -3.239411009 | 8.24E-05 |
| CCDC177 | -2.630404591 | 0.041006 | SDHC | -1.002044262 | 0.027255 |
| CCN1 | -2.929761604 | 1.42E-04 | SLC10A1 | -3.581084934 | 0.034808 |
| CCNG1 | -1.107965369 | 0.010103 | SLC11A2 | -1.66681981 | 0.002178 |
| CHRFAM7A | -4.0161074 | 0.022973 | SLC25A29 | -1.670204202 | 0.027255 |
| CPA5 | -4.28196816 | 5.28E-04 | SLC51A | -3.954189122 | 0.032323 |
| CST1 | -1.891030156 | 0.04684 | SPAG5-AS1 | -2.577385706 | 0.009805 |
| CTIF | -1.417093054 | 0.022103 | SPARCL1 | -2.675536403 | 0.011383 |
| DMBT1 | -1.477073181 | 0.032323 | SPTAN1 | -1.223386741 | 0.003172 |
| ECE1 | -1.183078638 | 0.010052 | ST6GAL1 | -1.770278643 | 0.020983 |
| ENSG00000257711 | -4.296522351 | 7.13E-04 | SURF4 | -1.369296066 | 0.009711 |
| GAL3ST1 | -6.125340052 | 1.69E-05 | TCF25 | -1.014783304 | 0.029827 |
| GLDC | -4.542739309 | 0.040334 | THBS1 | -1.835157091 | 0.001243 |
| GNA11 | -1.828080053 | 0.030157 | TRIM28 | -1.3383682 | 0.006946 |
| GRAMD1A | -1.033763767 | 0.00825 | TSPAN12 | -1.891732293 | 0.024727 |
| INO80 | -1.43237443 | 0.024317 | TTC7A | -1.086852135 | 0.045023 |
| ITGA5 | -1.328481127 | 0.002682 | UBN1 | -1.042632509 | 0.029827 |
| LHX5-AS1 | -3.317128213 | 0.016916 | UPF1 | -1.602580834 | 0.001578 |
| LINC01002 | -1.37347597 | 0.0393 | USP9Y | -3.256045785 | 0.043702 |
| LPCAT1 | -1.351951597 | 0.00904 | VPS9D1 | -1.768662732 | 0.002172 |
| LRPPRC | -1.379495137 | 0.044375 | ZER1 | -1.39856534 | 0.023607 |
| LSG1 | -1.744485678 | 0.027255 | | | |
| MKS1 | -2.862283185 | 0.001297 | | | |
| NDUFV1 | -1.460900005 | 0.045236 | | | |
| NF1 | -1.118275722 | 0.02813 | | | |
| NMRAL1 | -1.482657752 | 0.021664 | | | |
| NRK | -3.546189382 | 0.027255 | | | |
| OLFML2A | -2.198999808 | 0.029827 | | | |
| PCM1 | -1.167858084 | 0.024482 | | | |
| PDE4D | -1.528762915 | 0.021351 | | | |
Supplementary Table 4: Table of 71 differentially expressed genes downregulated in SSA+ samples in comparison to Control samples.

### Slide 18
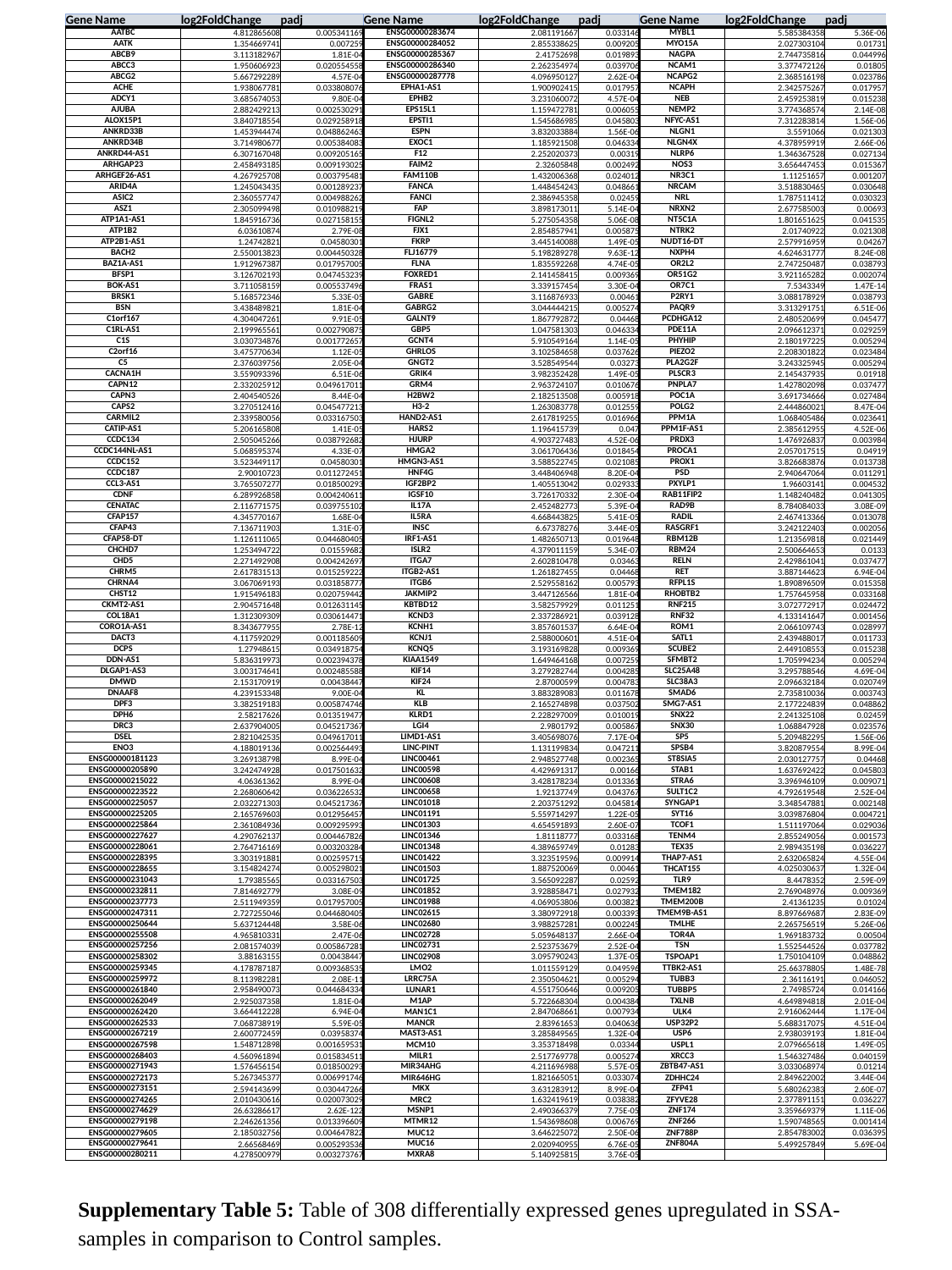

| Gene Name | log2FoldChange | padj | Gene Name | log2FoldChange | padj | Gene Name | log2FoldChange | padj |
| --- | --- | --- | --- | --- | --- | --- | --- | --- |
| AATBC | 4.812865608 | 0.005341169 | ENSG00000283674 | 2.081191667 | 0.033146 | MYBL1 | 5.585384358 | 5.36E-06 |
| AATK | 1.354669741 | 0.007259 | ENSG00000284052 | 2.855338625 | 0.009205 | MYO15A | 2.027303104 | 0.01731 |
| ABCB9 | 3.113182967 | 1.81E-04 | ENSG00000285367 | 2.41752698 | 0.019893 | NAGPA | 2.744735816 | 0.044996 |
| ABCC3 | 1.950606923 | 0.020554558 | ENSG00000286340 | 2.262354974 | 0.039706 | NCAM1 | 3.377472126 | 0.01805 |
| ABCG2 | 5.667292289 | 4.57E-04 | ENSG00000287778 | 4.096950127 | 2.62E-04 | NCAPG2 | 2.368516198 | 0.023786 |
| ACHE | 1.938067781 | 0.033808076 | EPHA1-AS1 | 1.900902415 | 0.017957 | NCAPH | 2.342575267 | 0.017957 |
| ADCY1 | 3.685674053 | 9.80E-04 | EPHB2 | 3.231060072 | 4.57E-04 | NEB | 2.459253819 | 0.015238 |
| AJUBA | 2.882429213 | 0.002530291 | EPS15L1 | 1.159472781 | 0.006055 | NEMP2 | 3.774368574 | 2.14E-08 |
| ALOX15P1 | 3.840718554 | 0.029258918 | EPSTI1 | 1.545686985 | 0.045803 | NFYC-AS1 | 7.312283814 | 1.56E-06 |
| ANKRD33B | 1.453944474 | 0.048862463 | ESPN | 3.832033884 | 1.56E-06 | NLGN1 | 3.5591066 | 0.021303 |
| ANKRD34B | 3.714980677 | 0.005384083 | EXOC1 | 1.185921508 | 0.046334 | NLGN4X | 4.378959919 | 2.66E-06 |
| ANKRD44-AS1 | 6.307167048 | 0.009205165 | F12 | 2.252020373 | 0.00319 | NLRP6 | 1.346367528 | 0.027134 |
| ARHGAP23 | 2.458493185 | 0.009193025 | FAIM2 | 2.32605848 | 0.002492 | NOS3 | 3.656447453 | 0.015367 |
| ARHGEF26-AS1 | 4.267925708 | 0.003795481 | FAM110B | 1.432006368 | 0.024012 | NR3C1 | 1.11251657 | 0.001207 |
| ARID4A | 1.245043435 | 0.001289237 | FANCA | 1.448454243 | 0.048661 | NRCAM | 3.518830465 | 0.030648 |
| ASIC2 | 2.360557747 | 0.004988262 | FANCI | 2.386945358 | 0.02459 | NRL | 1.787511412 | 0.030323 |
| ASZ1 | 2.305099498 | 0.010988219 | FAP | 3.898173011 | 5.14E-04 | NRXN2 | 2.677585003 | 0.00693 |
| ATP1A1-AS1 | 1.845916736 | 0.027158155 | FIGNL2 | 5.275054358 | 5.06E-08 | NT5C1A | 1.801651625 | 0.041535 |
| ATP1B2 | 6.03610874 | 2.79E-08 | FJX1 | 2.854857941 | 0.005875 | NTRK2 | 2.01740922 | 0.021308 |
| ATP2B1-AS1 | 1.24742821 | 0.04580301 | FKRP | 3.445140088 | 1.49E-05 | NUDT16-DT | 2.579916959 | 0.04267 |
| BACH2 | 2.550013823 | 0.004450328 | FLJ16779 | 5.198289278 | 9.63E-12 | NXPH4 | 4.624631777 | 8.24E-08 |
| BAZ1A-AS1 | 1.912967387 | 0.017957005 | FLNA | 1.835592268 | 4.74E-05 | OR2L2 | 2.747250487 | 0.038793 |
| BFSP1 | 3.126702193 | 0.047453239 | FOXRED1 | 2.141458415 | 0.009369 | OR51G2 | 3.921165282 | 0.002074 |
| BOK-AS1 | 3.711058159 | 0.005537496 | FRAS1 | 3.339157454 | 3.30E-04 | OR7C1 | 7.5343349 | 1.47E-14 |
| BRSK1 | 5.168572346 | 5.33E-05 | GABRE | 3.116876933 | 0.00461 | P2RY1 | 3.088178929 | 0.038793 |
| BSN | 3.438489821 | 1.81E-04 | GABRG2 | 3.044444215 | 0.005274 | PAQR9 | 3.313291751 | 6.51E-06 |
| C1orf167 | 4.304047261 | 9.91E-05 | GALNT9 | 1.867792872 | 0.04468 | PCDHGA12 | 2.480520699 | 0.045477 |
| C1RL-AS1 | 2.199965561 | 0.002790875 | GBP5 | 1.047581303 | 0.046334 | PDE11A | 2.096612371 | 0.029259 |
| C1S | 3.030734876 | 0.001772657 | GCNT4 | 5.910549164 | 1.14E-05 | PHYHIP | 2.180197225 | 0.005294 |
| C2orf16 | 3.475770634 | 1.12E-05 | GHRLOS | 3.102584658 | 0.037626 | PIEZO2 | 2.208301822 | 0.023484 |
| C5 | 2.376039756 | 2.05E-04 | GNGT2 | 3.528549544 | 0.03273 | PLA2G2F | 3.243325945 | 0.005294 |
| CACNA1H | 3.559093396 | 6.51E-06 | GRIK4 | 3.982352428 | 1.49E-05 | PLSCR3 | 2.145437935 | 0.01918 |
| CAPN12 | 2.332025912 | 0.049617011 | GRM4 | 2.963724107 | 0.010676 | PNPLA7 | 1.427802098 | 0.037477 |
| CAPN3 | 2.404540526 | 8.44E-04 | H2BW2 | 2.182513508 | 0.005918 | POC1A | 3.691734666 | 0.027484 |
| CAPS2 | 3.270512416 | 0.045477213 | H3-2 | 1.263083778 | 0.012559 | POLG2 | 2.444860021 | 8.47E-04 |
| CARMIL2 | 2.339580056 | 0.033167503 | HAND2-AS1 | 2.617819255 | 0.016966 | PPM1A | 1.068405486 | 0.023641 |
| CATIP-AS1 | 5.206165808 | 1.41E-05 | HARS2 | 1.196415739 | 0.047 | PPM1F-AS1 | 2.385612955 | 4.52E-06 |
| CCDC134 | 2.505045266 | 0.038792682 | HJURP | 4.903727483 | 4.52E-06 | PRDX3 | 1.476926837 | 0.003984 |
| CCDC144NL-AS1 | 5.068595374 | 4.33E-07 | HMGA2 | 3.061706436 | 0.018454 | PROCA1 | 2.057017515 | 0.04919 |
| CCDC152 | 3.523449117 | 0.04580301 | HMGN3-AS1 | 3.588522745 | 0.021085 | PROX1 | 3.826683876 | 0.013738 |
| CCDC187 | 2.90010723 | 0.011272451 | HNF4G | 3.448406948 | 8.20E-04 | PSD | 2.940647064 | 0.011291 |
| CCL3-AS1 | 3.765507277 | 0.018500293 | IGF2BP2 | 1.405513042 | 0.029333 | PXYLP1 | 1.96603141 | 0.004532 |
| CDNF | 6.289926858 | 0.004240611 | IGSF10 | 3.726170332 | 2.30E-04 | RAB11FIP2 | 1.148240482 | 0.041305 |
| CENATAC | 2.116771575 | 0.039755102 | IL17A | 2.452482773 | 5.39E-04 | RAD9B | 8.784084033 | 3.08E-09 |
| CFAP157 | 4.345770167 | 1.68E-04 | IL5RA | 4.668443825 | 5.41E-05 | RADIL | 2.467413366 | 0.013078 |
| CFAP43 | 7.136711903 | 1.31E-07 | INSC | 6.67378276 | 3.44E-05 | RASGRF1 | 3.242122403 | 0.002056 |
| CFAP58-DT | 1.126111065 | 0.044680405 | IRF1-AS1 | 1.482650713 | 0.019648 | RBM12B | 1.213569818 | 0.021449 |
| CHCHD7 | 1.253494722 | 0.01559682 | ISLR2 | 4.379011159 | 5.34E-07 | RBM24 | 2.500664653 | 0.0133 |
| CHD5 | 2.271492908 | 0.004242697 | ITGA7 | 2.602810478 | 0.03463 | RELN | 2.429861041 | 0.037477 |
| CHRM5 | 2.617831513 | 0.015259222 | ITGB2-AS1 | 1.261827455 | 0.04468 | RET | 3.887144623 | 6.94E-04 |
| CHRNA4 | 3.067069193 | 0.031858777 | ITGB6 | 2.529558162 | 0.005793 | RFPL1S | 1.890896509 | 0.015358 |
| CHST12 | 1.915496183 | 0.020759442 | JAKMIP2 | 3.447126566 | 1.81E-04 | RHOBTB2 | 1.757645958 | 0.033168 |
| CKMT2-AS1 | 2.904571648 | 0.012631145 | KBTBD12 | 3.582579929 | 0.011251 | RNF215 | 3.072772917 | 0.024472 |
| COL18A1 | 1.312309309 | 0.030614471 | KCND3 | 2.337286921 | 0.039128 | RNF32 | 4.133141647 | 0.001456 |
| CORO1A-AS1 | 8.343677955 | 2.78E-12 | KCNH1 | 3.857601537 | 6.64E-04 | ROM1 | 2.066109743 | 0.028997 |
| DACT3 | 4.117592029 | 0.001185609 | KCNJ1 | 2.588000601 | 4.51E-04 | SATL1 | 2.439488017 | 0.011733 |
| DCPS | 1.27948615 | 0.034918754 | KCNQ5 | 3.193169828 | 0.009369 | SCUBE2 | 2.449108553 | 0.015238 |
| DDN-AS1 | 5.836319973 | 0.002394378 | KIAA1549 | 1.649464168 | 0.007259 | SFMBT2 | 1.705994234 | 0.005294 |
| DLGAP1-AS3 | 3.003174641 | 0.002485588 | KIF14 | 3.279282744 | 0.004285 | SLC25A48 | 3.295788546 | 4.69E-04 |
| DMWD | 2.153170919 | 0.00438447 | KIF24 | 2.87000599 | 0.004783 | SLC38A3 | 2.096632184 | 0.020749 |
| DNAAF8 | 4.239153348 | 9.00E-04 | KL | 3.883289083 | 0.011678 | SMAD6 | 2.735810036 | 0.003743 |
| DPF3 | 3.382519183 | 0.005874746 | KLB | 2.165274898 | 0.037502 | SMG7-AS1 | 2.177224839 | 0.048862 |
| DPH6 | 2.58217626 | 0.013519477 | KLRD1 | 2.228297009 | 0.010019 | SNX22 | 2.241325108 | 0.02459 |
| DRC3 | 2.637904005 | 0.045217367 | LGI4 | 2.9801792 | 0.005867 | SNX30 | 1.068847928 | 0.023576 |
| DSEL | 2.821042535 | 0.049617011 | LIMD1-AS1 | 3.405698076 | 7.17E-04 | SP5 | 5.209482295 | 1.56E-06 |
| ENO3 | 4.188019136 | 0.002564493 | LINC-PINT | 1.131199834 | 0.047211 | SPSB4 | 3.820879554 | 8.99E-04 |
| ENSG00000181123 | 3.269138798 | 8.99E-04 | LINC00461 | 2.948527748 | 0.002365 | ST8SIA5 | 2.030127757 | 0.04468 |
| ENSG00000205890 | 3.242474928 | 0.017501632 | LINC00598 | 4.429691317 | 0.00166 | STAB1 | 1.637692422 | 0.045803 |
| ENSG00000215022 | 4.06361362 | 8.99E-04 | LINC00608 | 3.428178234 | 0.013361 | STRA6 | 3.396946109 | 0.009071 |
| ENSG00000223522 | 2.268060642 | 0.036226532 | LINC00658 | 1.92137749 | 0.043767 | SULT1C2 | 4.792619548 | 2.52E-04 |
| ENSG00000225057 | 2.032271303 | 0.045217367 | LINC01018 | 2.203751292 | 0.045814 | SYNGAP1 | 3.348547881 | 0.002148 |
| ENSG00000225205 | 2.165769603 | 0.012956457 | LINC01191 | 5.559714297 | 1.22E-05 | SYT16 | 3.039876804 | 0.004721 |
| ENSG00000225864 | 2.361084936 | 0.009295993 | LINC01303 | 4.654591893 | 2.60E-07 | TCOF1 | 1.511197064 | 0.029036 |
| ENSG00000227627 | 4.290762137 | 0.004467826 | LINC01346 | 1.81118777 | 0.033168 | TENM4 | 2.855249056 | 0.001573 |
| ENSG00000228061 | 2.764716169 | 0.003203284 | LINC01348 | 4.389659749 | 0.01283 | TEX35 | 2.989435198 | 0.036227 |
| ENSG00000228395 | 3.303191881 | 0.002595715 | LINC01422 | 3.323519596 | 0.009914 | THAP7-AS1 | 2.632065824 | 4.55E-04 |
| ENSG00000228655 | 3.154824274 | 0.005298021 | LINC01503 | 1.887520069 | 0.00461 | THCAT155 | 4.025030637 | 1.32E-04 |
| ENSG00000231043 | 1.79385565 | 0.033167503 | LINC01725 | 3.565092287 | 0.02592 | TLR9 | 8.4478352 | 2.59E-09 |
| ENSG00000232811 | 7.814692779 | 3.08E-09 | LINC01852 | 3.928858471 | 0.027932 | TMEM182 | 2.769048976 | 0.009369 |
| ENSG00000237773 | 2.511949359 | 0.017957005 | LINC01988 | 4.069053806 | 0.003821 | TMEM200B | 2.41361235 | 0.01024 |
| ENSG00000247311 | 2.727255046 | 0.044680405 | LINC02615 | 3.380972918 | 0.003393 | TMEM9B-AS1 | 8.897669687 | 2.83E-09 |
| ENSG00000250644 | 5.637124448 | 3.58E-06 | LINC02680 | 3.988257281 | 0.002245 | TMLHE | 2.265756519 | 5.26E-06 |
| ENSG00000255508 | 4.965810331 | 2.47E-06 | LINC02728 | 5.059648137 | 2.66E-04 | TOR4A | 1.969183732 | 0.00504 |
| ENSG00000257256 | 2.081574039 | 0.005867281 | LINC02731 | 2.523753679 | 2.52E-04 | TSN | 1.552544526 | 0.037782 |
| ENSG00000258302 | 3.88163155 | 0.00438447 | LINC02908 | 3.095790243 | 1.37E-05 | TSPOAP1 | 1.750104109 | 0.048862 |
| ENSG00000259345 | 4.178787187 | 0.009368535 | LMO2 | 1.011559129 | 0.049596 | TTBK2-AS1 | 25.66378805 | 1.48E-78 |
| ENSG00000259972 | 8.113982281 | 2.08E-11 | LRRC75A | 2.350504621 | 0.005294 | TUBB3 | 2.36116191 | 0.046052 |
| ENSG00000261840 | 2.958490073 | 0.044684334 | LUNAR1 | 4.551750646 | 0.009205 | TUBBP5 | 2.74985724 | 0.014166 |
| ENSG00000262049 | 2.925037358 | 1.81E-04 | M1AP | 5.722668304 | 0.004384 | TXLNB | 4.649894818 | 2.01E-04 |
| ENSG00000262420 | 3.664412228 | 6.94E-04 | MAN1C1 | 2.847068661 | 0.007934 | ULK4 | 2.916062444 | 1.17E-04 |
| ENSG00000262533 | 7.068738919 | 5.59E-05 | MANCR | 2.83961653 | 0.040636 | USP32P2 | 5.688317075 | 4.51E-04 |
| ENSG00000267219 | 2.600772459 | 0.03958374 | MAST3-AS1 | 3.285849565 | 1.32E-04 | USP6 | 2.938039193 | 1.81E-04 |
| ENSG00000267598 | 1.548712898 | 0.001659531 | MCM10 | 3.353718498 | 0.03344 | USPL1 | 2.079665618 | 1.49E-05 |
| ENSG00000268403 | 4.560961894 | 0.015834511 | MILR1 | 2.517769778 | 0.005274 | XRCC3 | 1.546327486 | 0.040159 |
| ENSG00000271943 | 1.576456154 | 0.018500293 | MIR34AHG | 4.211696988 | 5.57E-05 | ZBTB47-AS1 | 3.033068974 | 0.01214 |
| ENSG00000272173 | 5.267345377 | 0.006991746 | MIR646HG | 1.821665051 | 0.033074 | ZDHHC24 | 2.849622002 | 3.44E-04 |
| ENSG00000273151 | 2.594143699 | 0.030447266 | MKX | 3.631283912 | 8.99E-04 | ZFP41 | 5.680262383 | 2.60E-07 |
| ENSG00000274265 | 2.010430616 | 0.020073029 | MRC2 | 1.632419619 | 0.038382 | ZFYVE28 | 2.377891151 | 0.036227 |
| ENSG00000274629 | 26.63286617 | 2.62E-122 | MSNP1 | 2.490366379 | 7.75E-05 | ZNF174 | 3.359669379 | 1.11E-06 |
| ENSG00000279198 | 2.246261356 | 0.013396609 | MTMR12 | 1.543698608 | 0.006769 | ZNF266 | 1.590748565 | 0.001414 |
| ENSG00000279605 | 2.185032756 | 0.004647822 | MUC12 | 3.646225072 | 2.50E-06 | ZNF788P | 2.854783002 | 0.036395 |
| ENSG00000279641 | 2.66568469 | 0.005293536 | MUC16 | 2.020940955 | 6.76E-05 | ZNF804A | 5.499257849 | 5.69E-04 |
| ENSG00000280211 | 4.278500979 | 0.003273767 | MXRA8 | 5.140925815 | 3.76E-05 | | | |
Supplementary Table 5: Table of 308 differentially expressed genes upregulated in SSA- samples in comparison to Control samples.

### Slide 19
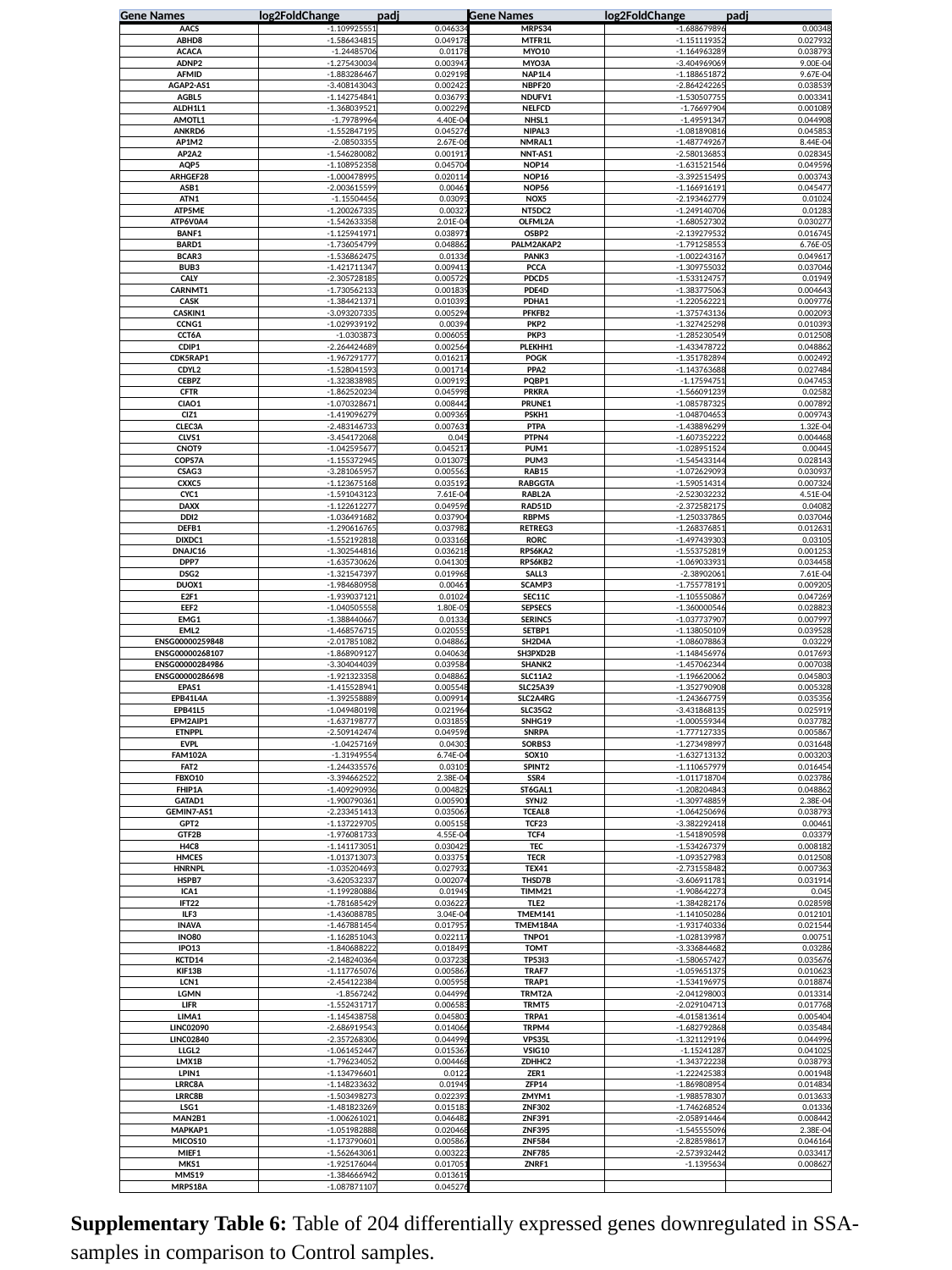

| Gene Names | log2FoldChange | padj | Gene Names | log2FoldChange | padj |
| --- | --- | --- | --- | --- | --- |
| AACS | -1.109925551 | 0.046334 | MRPS34 | -1.688679896 | 0.00348 |
| ABHD8 | -1.586434815 | 0.049178 | MTFR1L | -1.151119352 | 0.027932 |
| ACACA | -1.24485706 | 0.01178 | MYO10 | -1.164963289 | 0.038793 |
| ADNP2 | -1.275430034 | 0.003947 | MYO3A | -3.404969069 | 9.00E-04 |
| AFMID | -1.883286467 | 0.029198 | NAP1L4 | -1.188651872 | 9.67E-04 |
| AGAP2-AS1 | -3.408143043 | 0.002423 | NBPF20 | -2.864242265 | 0.038539 |
| AGBL5 | -1.142754841 | 0.036793 | NDUFV1 | -1.530507755 | 0.003341 |
| ALDH1L1 | -1.368039521 | 0.002296 | NELFCD | -1.76697904 | 0.001089 |
| AMOTL1 | -1.79789964 | 4.40E-04 | NHSL1 | -1.49591347 | 0.044908 |
| ANKRD6 | -1.552847195 | 0.045276 | NIPAL3 | -1.081890816 | 0.045853 |
| AP1M2 | -2.08503355 | 2.67E-06 | NMRAL1 | -1.487749267 | 8.44E-04 |
| AP2A2 | -1.546280082 | 0.001917 | NNT-AS1 | -2.580136853 | 0.028345 |
| AQP5 | -1.108952358 | 0.045704 | NOP14 | -1.631521546 | 0.049596 |
| ARHGEF28 | -1.000478995 | 0.020114 | NOP16 | -3.392515495 | 0.003743 |
| ASB1 | -2.003615599 | 0.00461 | NOP56 | -1.166916191 | 0.045477 |
| ATN1 | -1.15504456 | 0.03093 | NOX5 | -2.193462779 | 0.01024 |
| ATP5ME | -1.200267335 | 0.00327 | NT5DC2 | -1.249140706 | 0.01283 |
| ATP6V0A4 | -1.542633358 | 2.01E-04 | OLFML2A | -1.680527302 | 0.030277 |
| BANF1 | -1.125941971 | 0.038971 | OSBP2 | -2.139279532 | 0.016745 |
| BARD1 | -1.736054799 | 0.048862 | PALM2AKAP2 | -1.791258553 | 6.76E-05 |
| BCAR3 | -1.536862475 | 0.01336 | PANK3 | -1.002243167 | 0.049617 |
| BUB3 | -1.421711347 | 0.009413 | PCCA | -1.309755032 | 0.037046 |
| CALY | -2.305728185 | 0.005729 | PDCD5 | -1.533124757 | 0.01949 |
| CARNMT1 | -1.730562133 | 0.001839 | PDE4D | -1.383775063 | 0.004643 |
| CASK | -1.384421371 | 0.010393 | PDHA1 | -1.220562221 | 0.009776 |
| CASKIN1 | -3.093207335 | 0.005294 | PFKFB2 | -1.375743136 | 0.002093 |
| CCNG1 | -1.029939192 | 0.00394 | PKP2 | -1.327425298 | 0.010393 |
| CCT6A | -1.0303873 | 0.006055 | PKP3 | -1.285230549 | 0.012508 |
| CDIP1 | -2.264424689 | 0.002564 | PLEKHH1 | -1.433478722 | 0.048862 |
| CDK5RAP1 | -1.967291777 | 0.016217 | POGK | -1.351782894 | 0.002492 |
| CDYL2 | -1.528041593 | 0.001714 | PPA2 | -1.143763688 | 0.027484 |
| CEBPZ | -1.323838985 | 0.009193 | PQBP1 | -1.17594751 | 0.047453 |
| CFTR | -1.862520234 | 0.045998 | PRKRA | -1.566091239 | 0.02582 |
| CIAO1 | -1.070328671 | 0.008442 | PRUNE1 | -1.085787325 | 0.007892 |
| CIZ1 | -1.419096279 | 0.009369 | PSKH1 | -1.048704653 | 0.009743 |
| CLEC3A | -2.483146733 | 0.007631 | PTPA | -1.438896299 | 1.32E-04 |
| CLVS1 | -3.454172068 | 0.045 | PTPN4 | -1.607352222 | 0.004468 |
| CNOT9 | -1.042595677 | 0.045217 | PUM1 | -1.028951524 | 0.00445 |
| COPS7A | -1.155372945 | 0.013075 | PUM3 | -1.545433144 | 0.028143 |
| CSAG3 | -3.281065957 | 0.005563 | RAB15 | -1.072629093 | 0.030937 |
| CXXC5 | -1.123675168 | 0.035192 | RABGGTA | -1.590514314 | 0.007324 |
| CYC1 | -1.591043123 | 7.61E-04 | RABL2A | -2.523032232 | 4.51E-04 |
| DAXX | -1.122612277 | 0.049596 | RAD51D | -2.372582175 | 0.04082 |
| DDI2 | -1.036491682 | 0.037904 | RBPMS | -1.250337865 | 0.037046 |
| DEFB1 | -1.290616765 | 0.037982 | RETREG3 | -1.268376851 | 0.012631 |
| DIXDC1 | -1.552192818 | 0.033168 | RORC | -1.497439303 | 0.03105 |
| DNAJC16 | -1.302544816 | 0.036218 | RPS6KA2 | -1.553752819 | 0.001253 |
| DPP7 | -1.635730626 | 0.041305 | RPS6KB2 | -1.069033931 | 0.034458 |
| DSG2 | -1.321547397 | 0.019968 | SALL3 | -2.38902061 | 7.61E-04 |
| DUOX1 | -1.984680958 | 0.00461 | SCAMP3 | -1.755778191 | 0.009205 |
| E2F1 | -1.939037121 | 0.01024 | SEC11C | -1.105550867 | 0.047269 |
| EEF2 | -1.040505558 | 1.80E-05 | SEPSECS | -1.360000546 | 0.028823 |
| EMG1 | -1.388440667 | 0.01336 | SERINC5 | -1.037737907 | 0.007997 |
| EML2 | -1.468576715 | 0.020555 | SETBP1 | -1.138050109 | 0.039528 |
| ENSG00000259848 | -2.017851082 | 0.048862 | SH2D4A | -1.086078863 | 0.03229 |
| ENSG00000268107 | -1.868909127 | 0.040636 | SH3PXD2B | -1.148456976 | 0.017693 |
| ENSG00000284986 | -3.304044039 | 0.039584 | SHANK2 | -1.457062344 | 0.007038 |
| ENSG00000286698 | -1.921323358 | 0.048862 | SLC11A2 | -1.196620062 | 0.045803 |
| EPAS1 | -1.415528941 | 0.005548 | SLC25A39 | -1.352790908 | 0.005328 |
| EPB41L4A | -1.392558889 | 0.009914 | SLC2A4RG | -1.243667759 | 0.035356 |
| EPB41L5 | -1.049480198 | 0.021964 | SLC35G2 | -3.431868135 | 0.025919 |
| EPM2AIP1 | -1.637198777 | 0.031859 | SNHG19 | -1.000559344 | 0.037782 |
| ETNPPL | -2.509142474 | 0.049596 | SNRPA | -1.777127335 | 0.005867 |
| EVPL | -1.04257169 | 0.04303 | SORBS3 | -1.273498997 | 0.031648 |
| FAM102A | -1.31949554 | 6.74E-04 | SOX10 | -1.632713132 | 0.003203 |
| FAT2 | -1.244335576 | 0.03105 | SPINT2 | -1.110657979 | 0.016454 |
| FBXO10 | -3.394662522 | 2.38E-04 | SSR4 | -1.011718704 | 0.023786 |
| FHIP1A | -1.409290936 | 0.004829 | ST6GAL1 | -1.208204843 | 0.048862 |
| GATAD1 | -1.900790361 | 0.005901 | SYNJ2 | -1.309748859 | 2.38E-04 |
| GEMIN7-AS1 | -2.233451413 | 0.035067 | TCEAL8 | -1.064250696 | 0.038793 |
| GPT2 | -1.137229705 | 0.005158 | TCF23 | -3.382292418 | 0.00461 |
| GTF2B | -1.976081733 | 4.55E-04 | TCF4 | -1.541890598 | 0.03379 |
| H4C8 | -1.141173051 | 0.030425 | TEC | -1.534267379 | 0.008182 |
| HMCES | -1.013713073 | 0.033751 | TECR | -1.093527983 | 0.012508 |
| HNRNPL | -1.035204693 | 0.027932 | TEX41 | -2.731558482 | 0.007363 |
| HSPB7 | -3.620532337 | 0.002074 | THSD7B | -3.606911781 | 0.031914 |
| ICA1 | -1.199280886 | 0.01949 | TIMM21 | -1.908642273 | 0.045 |
| IFT22 | -1.781685429 | 0.036227 | TLE2 | -1.384282176 | 0.028598 |
| ILF3 | -1.436088785 | 3.04E-04 | TMEM141 | -1.141050286 | 0.012101 |
| INAVA | -1.467881454 | 0.017957 | TMEM184A | -1.931740336 | 0.021544 |
| INO80 | -1.162851043 | 0.022117 | TNPO1 | -1.028139987 | 0.00751 |
| IPO13 | -1.840688222 | 0.018495 | TOMT | -3.336844682 | 0.03286 |
| KCTD14 | -2.148240364 | 0.037238 | TP53I3 | -1.580657427 | 0.035676 |
| KIF13B | -1.117765076 | 0.005867 | TRAF7 | -1.059651375 | 0.010623 |
| LCN1 | -2.454122384 | 0.005958 | TRAP1 | -1.534196975 | 0.018874 |
| LGMN | -1.8567242 | 0.044996 | TRMT2A | -2.041298003 | 0.013314 |
| LIFR | -1.552431717 | 0.006583 | TRMT5 | -2.029104713 | 0.017768 |
| LIMA1 | -1.145438758 | 0.045803 | TRPA1 | -4.015813614 | 0.005404 |
| LINC02090 | -2.686919543 | 0.014066 | TRPM4 | -1.682792868 | 0.035484 |
| LINC02840 | -2.357268306 | 0.044996 | VPS35L | -1.321129196 | 0.044996 |
| LLGL2 | -1.061452447 | 0.015367 | VSIG10 | -1.15241287 | 0.041025 |
| LMX1B | -1.796234052 | 0.004468 | ZDHHC2 | -1.343722238 | 0.038793 |
| LPIN1 | -1.134796601 | 0.0122 | ZER1 | -1.222425383 | 0.001948 |
| LRRC8A | -1.148233632 | 0.01949 | ZFP14 | -1.869808954 | 0.014834 |
| LRRC8B | -1.503498273 | 0.022393 | ZMYM1 | -1.988578307 | 0.013633 |
| LSG1 | -1.481823269 | 0.015183 | ZNF302 | -1.746268524 | 0.01336 |
| MAN2B1 | -1.006261021 | 0.046482 | ZNF391 | -2.058914464 | 0.008442 |
| MAPKAP1 | -1.051982888 | 0.020468 | ZNF395 | -1.545555096 | 2.38E-04 |
| MICOS10 | -1.173790601 | 0.005867 | ZNF584 | -2.828598617 | 0.046164 |
| MIEF1 | -1.562643061 | 0.003223 | ZNF785 | -2.573932442 | 0.033417 |
| MKS1 | -1.925176044 | 0.017051 | ZNRF1 | -1.1395634 | 0.008627 |
| MMS19 | -1.384666942 | 0.013619 | | | |
| MRPS18A | -1.087871107 | 0.045276 | | | |
Supplementary Table 6: Table of 204 differentially expressed genes downregulated in SSA- samples in comparison to Control samples.

### Slide 20
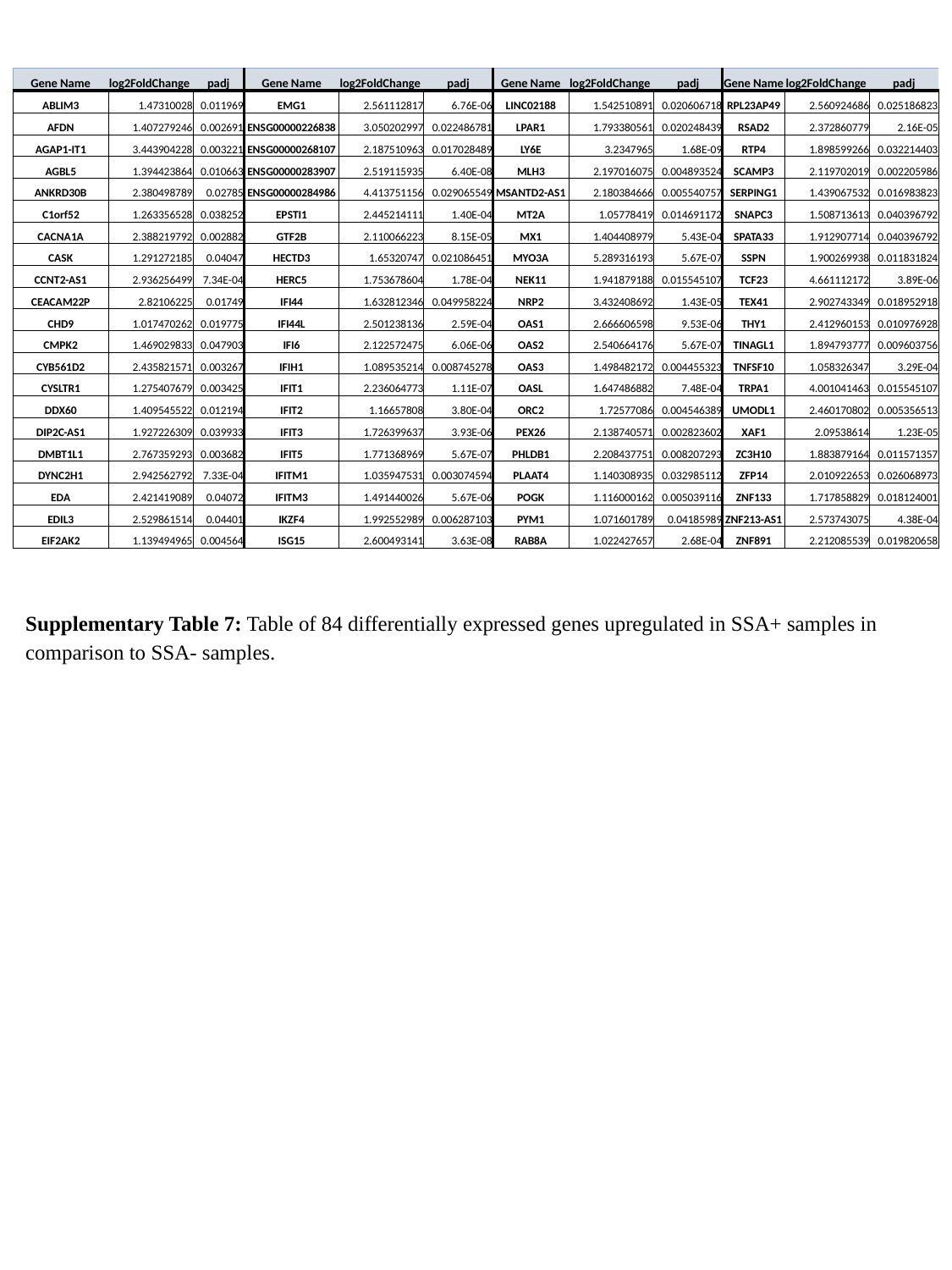

| Gene Name | log2FoldChange | padj | Gene Name | log2FoldChange | padj | Gene Name | log2FoldChange | padj | Gene Name | log2FoldChange | padj |
| --- | --- | --- | --- | --- | --- | --- | --- | --- | --- | --- | --- |
| ABLIM3 | 1.47310028 | 0.011969 | EMG1 | 2.561112817 | 6.76E-06 | LINC02188 | 1.542510891 | 0.020606718 | RPL23AP49 | 2.560924686 | 0.025186823 |
| AFDN | 1.407279246 | 0.002691 | ENSG00000226838 | 3.050202997 | 0.022486781 | LPAR1 | 1.793380561 | 0.020248439 | RSAD2 | 2.372860779 | 2.16E-05 |
| AGAP1-IT1 | 3.443904228 | 0.003221 | ENSG00000268107 | 2.187510963 | 0.017028489 | LY6E | 3.2347965 | 1.68E-09 | RTP4 | 1.898599266 | 0.032214403 |
| AGBL5 | 1.394423864 | 0.010663 | ENSG00000283907 | 2.519115935 | 6.40E-08 | MLH3 | 2.197016075 | 0.004893524 | SCAMP3 | 2.119702019 | 0.002205986 |
| ANKRD30B | 2.380498789 | 0.02785 | ENSG00000284986 | 4.413751156 | 0.029065549 | MSANTD2-AS1 | 2.180384666 | 0.005540757 | SERPING1 | 1.439067532 | 0.016983823 |
| C1orf52 | 1.263356528 | 0.038252 | EPSTI1 | 2.445214111 | 1.40E-04 | MT2A | 1.05778419 | 0.014691172 | SNAPC3 | 1.508713613 | 0.040396792 |
| CACNA1A | 2.388219792 | 0.002882 | GTF2B | 2.110066223 | 8.15E-05 | MX1 | 1.404408979 | 5.43E-04 | SPATA33 | 1.912907714 | 0.040396792 |
| CASK | 1.291272185 | 0.04047 | HECTD3 | 1.65320747 | 0.021086451 | MYO3A | 5.289316193 | 5.67E-07 | SSPN | 1.900269938 | 0.011831824 |
| CCNT2-AS1 | 2.936256499 | 7.34E-04 | HERC5 | 1.753678604 | 1.78E-04 | NEK11 | 1.941879188 | 0.015545107 | TCF23 | 4.661112172 | 3.89E-06 |
| CEACAM22P | 2.82106225 | 0.01749 | IFI44 | 1.632812346 | 0.049958224 | NRP2 | 3.432408692 | 1.43E-05 | TEX41 | 2.902743349 | 0.018952918 |
| CHD9 | 1.017470262 | 0.019775 | IFI44L | 2.501238136 | 2.59E-04 | OAS1 | 2.666606598 | 9.53E-06 | THY1 | 2.412960153 | 0.010976928 |
| CMPK2 | 1.469029833 | 0.047903 | IFI6 | 2.122572475 | 6.06E-06 | OAS2 | 2.540664176 | 5.67E-07 | TINAGL1 | 1.894793777 | 0.009603756 |
| CYB561D2 | 2.435821571 | 0.003267 | IFIH1 | 1.089535214 | 0.008745278 | OAS3 | 1.498482172 | 0.004455323 | TNFSF10 | 1.058326347 | 3.29E-04 |
| CYSLTR1 | 1.275407679 | 0.003425 | IFIT1 | 2.236064773 | 1.11E-07 | OASL | 1.647486882 | 7.48E-04 | TRPA1 | 4.001041463 | 0.015545107 |
| DDX60 | 1.409545522 | 0.012194 | IFIT2 | 1.16657808 | 3.80E-04 | ORC2 | 1.72577086 | 0.004546389 | UMODL1 | 2.460170802 | 0.005356513 |
| DIP2C-AS1 | 1.927226309 | 0.039933 | IFIT3 | 1.726399637 | 3.93E-06 | PEX26 | 2.138740571 | 0.002823602 | XAF1 | 2.09538614 | 1.23E-05 |
| DMBT1L1 | 2.767359293 | 0.003682 | IFIT5 | 1.771368969 | 5.67E-07 | PHLDB1 | 2.208437751 | 0.008207293 | ZC3H10 | 1.883879164 | 0.011571357 |
| DYNC2H1 | 2.942562792 | 7.33E-04 | IFITM1 | 1.035947531 | 0.003074594 | PLAAT4 | 1.140308935 | 0.032985112 | ZFP14 | 2.010922653 | 0.026068973 |
| EDA | 2.421419089 | 0.04072 | IFITM3 | 1.491440026 | 5.67E-06 | POGK | 1.116000162 | 0.005039116 | ZNF133 | 1.717858829 | 0.018124001 |
| EDIL3 | 2.529861514 | 0.04401 | IKZF4 | 1.992552989 | 0.006287103 | PYM1 | 1.071601789 | 0.04185989 | ZNF213-AS1 | 2.573743075 | 4.38E-04 |
| EIF2AK2 | 1.139494965 | 0.004564 | ISG15 | 2.600493141 | 3.63E-08 | RAB8A | 1.022427657 | 2.68E-04 | ZNF891 | 2.212085539 | 0.019820658 |
Supplementary Table 7: Table of 84 differentially expressed genes upregulated in SSA+ samples in comparison to SSA- samples.

### Slide 21
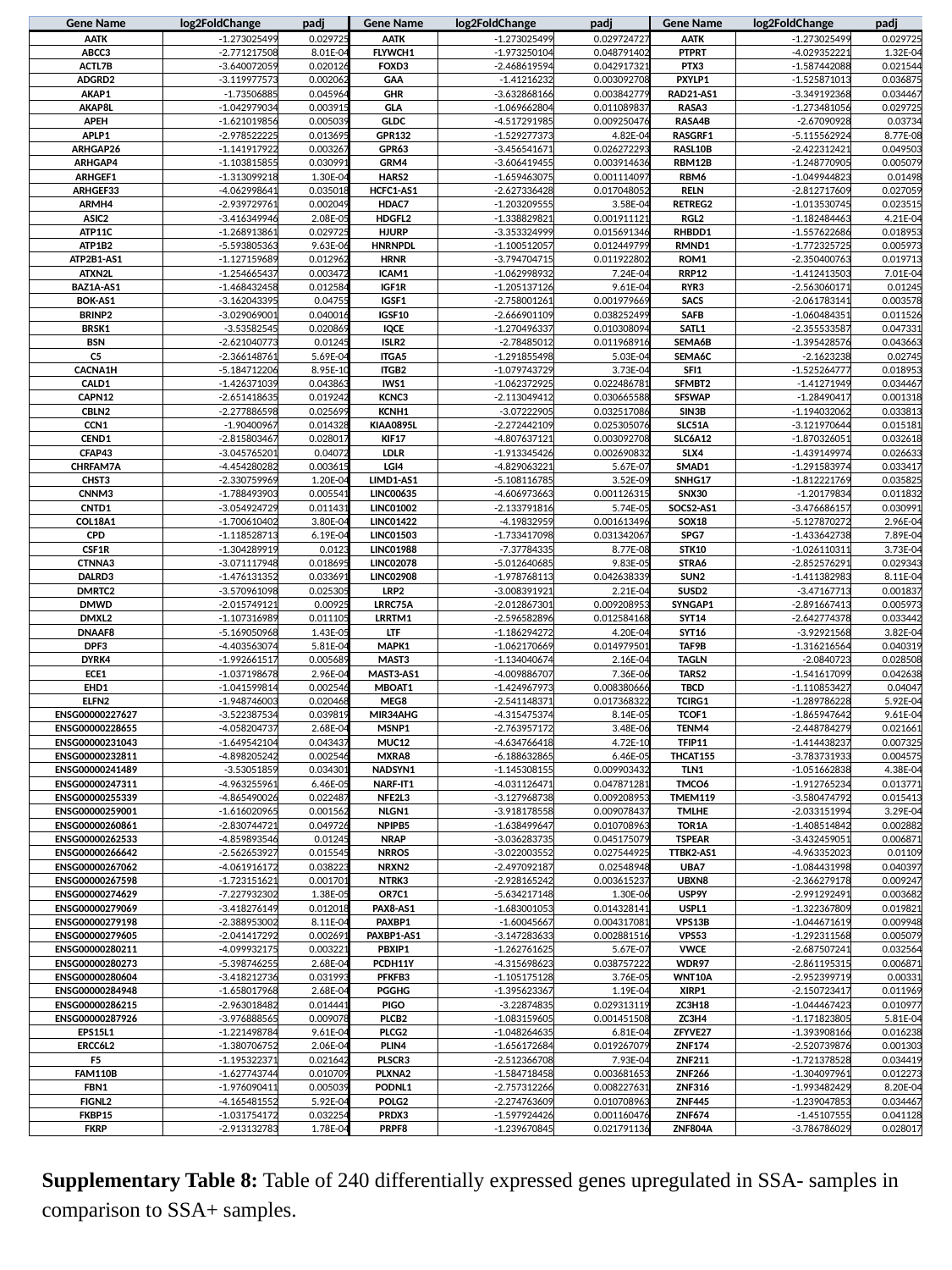

| Gene Name | log2FoldChange | padj | Gene Name | log2FoldChange | padj | Gene Name | log2FoldChange | padj |
| --- | --- | --- | --- | --- | --- | --- | --- | --- |
| AATK | -1.273025499 | 0.029725 | AATK | -1.273025499 | 0.029724727 | AATK | -1.273025499 | 0.029725 |
| ABCC3 | -2.771217508 | 8.01E-04 | FLYWCH1 | -1.973250104 | 0.048791402 | PTPRT | -4.029352221 | 1.32E-04 |
| ACTL7B | -3.640072059 | 0.020126 | FOXD3 | -2.468619594 | 0.042917321 | PTX3 | -1.587442088 | 0.021544 |
| ADGRD2 | -3.119977573 | 0.002062 | GAA | -1.41216232 | 0.003092708 | PXYLP1 | -1.525871013 | 0.036875 |
| AKAP1 | -1.73506885 | 0.045964 | GHR | -3.632868166 | 0.003842779 | RAD21-AS1 | -3.349192368 | 0.034467 |
| AKAP8L | -1.042979034 | 0.003915 | GLA | -1.069662804 | 0.011089837 | RASA3 | -1.273481056 | 0.029725 |
| APEH | -1.621019856 | 0.005039 | GLDC | -4.517291985 | 0.009250476 | RASA4B | -2.67090928 | 0.03734 |
| APLP1 | -2.978522225 | 0.013695 | GPR132 | -1.529277373 | 4.82E-04 | RASGRF1 | -5.115562924 | 8.77E-08 |
| ARHGAP26 | -1.141917922 | 0.003267 | GPR63 | -3.456541671 | 0.026272293 | RASL10B | -2.422312421 | 0.049503 |
| ARHGAP4 | -1.103815855 | 0.030991 | GRM4 | -3.606419455 | 0.003914636 | RBM12B | -1.248770905 | 0.005079 |
| ARHGEF1 | -1.313099218 | 1.30E-04 | HARS2 | -1.659463075 | 0.001114097 | RBM6 | -1.049944823 | 0.01498 |
| ARHGEF33 | -4.062998641 | 0.035018 | HCFC1-AS1 | -2.627336428 | 0.017048052 | RELN | -2.812717609 | 0.027059 |
| ARMH4 | -2.939729761 | 0.002049 | HDAC7 | -1.203209555 | 3.58E-04 | RETREG2 | -1.013530745 | 0.023515 |
| ASIC2 | -3.416349946 | 2.08E-05 | HDGFL2 | -1.338829821 | 0.001911121 | RGL2 | -1.182484463 | 4.21E-04 |
| ATP11C | -1.268913861 | 0.029725 | HJURP | -3.353324999 | 0.015691346 | RHBDD1 | -1.557622686 | 0.018953 |
| ATP1B2 | -5.593805363 | 9.63E-06 | HNRNPDL | -1.100512057 | 0.012449799 | RMND1 | -1.772325725 | 0.005973 |
| ATP2B1-AS1 | -1.127159689 | 0.012962 | HRNR | -3.794704715 | 0.011922802 | ROM1 | -2.350400763 | 0.019713 |
| ATXN2L | -1.254665437 | 0.003472 | ICAM1 | -1.062998932 | 7.24E-04 | RRP12 | -1.412413503 | 7.01E-04 |
| BAZ1A-AS1 | -1.468432458 | 0.012584 | IGF1R | -1.205137126 | 9.61E-04 | RYR3 | -2.563060171 | 0.01245 |
| BOK-AS1 | -3.162043395 | 0.04755 | IGSF1 | -2.758001261 | 0.001979669 | SACS | -2.061783141 | 0.003578 |
| BRINP2 | -3.029069001 | 0.040016 | IGSF10 | -2.666901109 | 0.038252499 | SAFB | -1.060484351 | 0.011526 |
| BRSK1 | -3.53582545 | 0.020869 | IQCE | -1.270496337 | 0.010308094 | SATL1 | -2.355533587 | 0.047331 |
| BSN | -2.621040773 | 0.01245 | ISLR2 | -2.78485012 | 0.011968916 | SEMA6B | -1.395428576 | 0.043663 |
| C5 | -2.366148761 | 5.69E-04 | ITGA5 | -1.291855498 | 5.03E-04 | SEMA6C | -2.1623238 | 0.02745 |
| CACNA1H | -5.184712206 | 8.95E-10 | ITGB2 | -1.079743729 | 3.73E-04 | SFI1 | -1.525264777 | 0.018953 |
| CALD1 | -1.426371039 | 0.043863 | IWS1 | -1.062372925 | 0.022486781 | SFMBT2 | -1.41271949 | 0.034467 |
| CAPN12 | -2.651418635 | 0.019242 | KCNC3 | -2.113049412 | 0.030665588 | SFSWAP | -1.28490417 | 0.001318 |
| CBLN2 | -2.277886598 | 0.025699 | KCNH1 | -3.07222905 | 0.032517086 | SIN3B | -1.194032062 | 0.033813 |
| CCN1 | -1.90400967 | 0.014328 | KIAA0895L | -2.272442109 | 0.025305076 | SLC51A | -3.121970644 | 0.015181 |
| CEND1 | -2.815803467 | 0.028017 | KIF17 | -4.807637121 | 0.003092708 | SLC6A12 | -1.870326051 | 0.032618 |
| CFAP43 | -3.045765201 | 0.04072 | LDLR | -1.913345426 | 0.002690832 | SLX4 | -1.439149974 | 0.026633 |
| CHRFAM7A | -4.454280282 | 0.003615 | LGI4 | -4.829063221 | 5.67E-07 | SMAD1 | -1.291583974 | 0.033417 |
| CHST3 | -2.330759969 | 1.20E-04 | LIMD1-AS1 | -5.108116785 | 3.52E-09 | SNHG17 | -1.812221769 | 0.035825 |
| CNNM3 | -1.788493903 | 0.005541 | LINC00635 | -4.606973663 | 0.001126315 | SNX30 | -1.20179834 | 0.011832 |
| CNTD1 | -3.054924729 | 0.011431 | LINC01002 | -2.133791816 | 5.74E-05 | SOCS2-AS1 | -3.476686157 | 0.030991 |
| COL18A1 | -1.700610402 | 3.80E-04 | LINC01422 | -4.19832959 | 0.001613496 | SOX18 | -5.127870272 | 2.96E-04 |
| CPD | -1.118528713 | 6.19E-04 | LINC01503 | -1.733417098 | 0.031342067 | SPG7 | -1.433642738 | 7.89E-04 |
| CSF1R | -1.304289919 | 0.0123 | LINC01988 | -7.37784335 | 8.77E-08 | STK10 | -1.026110311 | 3.73E-04 |
| CTNNA3 | -3.071117948 | 0.018695 | LINC02078 | -5.012640685 | 9.83E-05 | STRA6 | -2.852576291 | 0.029343 |
| DALRD3 | -1.476131352 | 0.033691 | LINC02908 | -1.978768113 | 0.042638339 | SUN2 | -1.411382983 | 8.11E-04 |
| DMRTC2 | -3.570961098 | 0.025305 | LRP2 | -3.008391921 | 2.21E-04 | SUSD2 | -3.47167713 | 0.001837 |
| DMWD | -2.015749121 | 0.00925 | LRRC75A | -2.012867301 | 0.009208953 | SYNGAP1 | -2.891667413 | 0.005973 |
| DMXL2 | -1.107316989 | 0.011105 | LRRTM1 | -2.596582896 | 0.012584168 | SYT14 | -2.642774378 | 0.033442 |
| DNAAF8 | -5.169050968 | 1.43E-05 | LTF | -1.186294272 | 4.20E-04 | SYT16 | -3.92921568 | 3.82E-04 |
| DPF3 | -4.403563074 | 5.81E-04 | MAPK1 | -1.062170669 | 0.014979501 | TAF9B | -1.316216564 | 0.040319 |
| DYRK4 | -1.992661517 | 0.005689 | MAST3 | -1.134040674 | 2.16E-04 | TAGLN | -2.0840723 | 0.028508 |
| ECE1 | -1.037198678 | 2.96E-04 | MAST3-AS1 | -4.009886707 | 7.36E-06 | TARS2 | -1.541617099 | 0.042638 |
| EHD1 | -1.041599814 | 0.002546 | MBOAT1 | -1.424967973 | 0.008380666 | TBCD | -1.110853427 | 0.04047 |
| ELFN2 | -1.948746003 | 0.020468 | MEG8 | -2.541148371 | 0.017368322 | TCIRG1 | -1.289786228 | 5.92E-04 |
| ENSG00000227627 | -3.522387534 | 0.039819 | MIR34AHG | -4.315475374 | 8.14E-05 | TCOF1 | -1.865947642 | 9.61E-04 |
| ENSG00000228655 | -4.058204737 | 2.68E-04 | MSNP1 | -2.763957172 | 3.48E-06 | TENM4 | -2.448784279 | 0.021661 |
| ENSG00000231043 | -1.649542104 | 0.043437 | MUC12 | -4.634766418 | 4.72E-10 | TFIP11 | -1.414438237 | 0.007325 |
| ENSG00000232811 | -4.898205242 | 0.002546 | MXRA8 | -6.188632865 | 6.46E-05 | THCAT155 | -3.783731933 | 0.004575 |
| ENSG00000241489 | -3.53051859 | 0.034301 | NADSYN1 | -1.145308155 | 0.009903432 | TLN1 | -1.051662838 | 4.38E-04 |
| ENSG00000247311 | -4.963255961 | 6.46E-05 | NARF-IT1 | -4.031126471 | 0.047871281 | TMCO6 | -1.912765234 | 0.013771 |
| ENSG00000255339 | -4.865490026 | 0.022487 | NFE2L3 | -3.127968738 | 0.009208953 | TMEM119 | -3.580474792 | 0.015413 |
| ENSG00000259001 | -1.616020965 | 0.001562 | NLGN1 | -3.918178558 | 0.009078437 | TMLHE | -2.033151994 | 3.29E-04 |
| ENSG00000260861 | -2.830744721 | 0.049726 | NPIPB5 | -1.638499647 | 0.010708963 | TOR1A | -1.408514842 | 0.002882 |
| ENSG00000262533 | -4.859893546 | 0.01245 | NRAP | -3.036283735 | 0.045175079 | TSPEAR | -3.432459051 | 0.006871 |
| ENSG00000266642 | -2.562653927 | 0.015545 | NRROS | -3.022003552 | 0.027544925 | TTBK2-AS1 | -4.963352023 | 0.01109 |
| ENSG00000267062 | -4.061916172 | 0.038223 | NRXN2 | -2.497092187 | 0.02548948 | UBA7 | -1.084431998 | 0.040397 |
| ENSG00000267598 | -1.723151621 | 0.001701 | NTRK3 | -2.928165242 | 0.003615237 | UBXN8 | -2.366279178 | 0.009247 |
| ENSG00000274629 | -7.227932302 | 1.38E-05 | OR7C1 | -5.634217148 | 1.30E-06 | USP9Y | -2.991292491 | 0.003682 |
| ENSG00000279069 | -3.418276149 | 0.012018 | PAX8-AS1 | -1.683001053 | 0.014328141 | USPL1 | -1.322367809 | 0.019821 |
| ENSG00000279198 | -2.388953002 | 8.11E-04 | PAXBP1 | -1.60045667 | 0.004317081 | VPS13B | -1.044671619 | 0.009948 |
| ENSG00000279605 | -2.041417292 | 0.002691 | PAXBP1-AS1 | -3.147283633 | 0.002881516 | VPS53 | -1.292311568 | 0.005079 |
| ENSG00000280211 | -4.099932175 | 0.003221 | PBXIP1 | -1.262761625 | 5.67E-07 | VWCE | -2.687507241 | 0.032564 |
| ENSG00000280273 | -5.398746255 | 2.68E-04 | PCDH11Y | -4.315698623 | 0.038757222 | WDR97 | -2.861195315 | 0.006871 |
| ENSG00000280604 | -3.418212736 | 0.031993 | PFKFB3 | -1.105175128 | 3.76E-05 | WNT10A | -2.952399719 | 0.00331 |
| ENSG00000284948 | -1.658017968 | 2.68E-04 | PGGHG | -1.395623367 | 1.19E-04 | XIRP1 | -2.150723417 | 0.011969 |
| ENSG00000286215 | -2.963018482 | 0.014441 | PIGO | -3.22874835 | 0.029313119 | ZC3H18 | -1.044467423 | 0.010977 |
| ENSG00000287926 | -3.976888565 | 0.009078 | PLCB2 | -1.083159605 | 0.001451508 | ZC3H4 | -1.171823805 | 5.81E-04 |
| EPS15L1 | -1.221498784 | 9.61E-04 | PLCG2 | -1.048264635 | 6.81E-04 | ZFYVE27 | -1.393908166 | 0.016238 |
| ERCC6L2 | -1.380706752 | 2.06E-04 | PLIN4 | -1.656172684 | 0.019267079 | ZNF174 | -2.520739876 | 0.001303 |
| F5 | -1.195322371 | 0.021642 | PLSCR3 | -2.512366708 | 7.93E-04 | ZNF211 | -1.721378528 | 0.034419 |
| FAM110B | -1.627743744 | 0.010709 | PLXNA2 | -1.584718458 | 0.003681653 | ZNF266 | -1.304097961 | 0.012273 |
| FBN1 | -1.976090411 | 0.005039 | PODNL1 | -2.757312266 | 0.008227631 | ZNF316 | -1.993482429 | 8.20E-04 |
| FIGNL2 | -4.165481552 | 5.92E-04 | POLG2 | -2.274763609 | 0.010708963 | ZNF445 | -1.239047853 | 0.034467 |
| FKBP15 | -1.031754172 | 0.032254 | PRDX3 | -1.597924426 | 0.001160476 | ZNF674 | -1.45107555 | 0.041128 |
| FKRP | -2.913132783 | 1.78E-04 | PRPF8 | -1.239670845 | 0.021791136 | ZNF804A | -3.786786029 | 0.028017 |
Supplementary Table 8: Table of 240 differentially expressed genes upregulated in SSA- samples in comparison to SSA+ samples.
